# Arginine-vasopressin mediates counter-regulatory glucagon release and is diminished in type 1 diabetes

**DOI:** 10.1101/2020.01.30.927426

**Authors:** Angela Kim, Jakob G. Knudsen, Joseph C. Madara, Anna Benrick, Thomas Hill, Lina Abdul Kadir, Joely A. Kellard, Lisa Mellander, Caroline Miranda, Haopeng Lin, Timothy James, Kinga Suba, Aliya F. Spigelman, Yanling Wu, Patrick E. MacDonald, Ingrid Wernstedt Asterholm, Tore Magnussen, Mikkel Christensen, Tina Visboll, Victoria Salem, Filip K. Knop, Patrik Rorsman, Bradford B. Lowell, Linford J.B. Briant

## Abstract

Insulin-induced hypoglycemia is a major barrier to the treatment of type-1 diabetes (T1D). Accordingly, it is important that we understand the mechanisms regulating the circulating levels of glucagon – the body’s principal blood glucose-elevating hormone which is secreted from alpha-cells of the pancreatic islets. Varying glucose over the range of concentrations that occur physiologically between the fed and fuel-deprived states (from 8 to 4 mM) has no significant effect on glucagon secretion in isolated islets (*in vitro*) and yet associates with dramatic changes in plasma glucagon *in vivo*. The identity of the systemic factor(s) that stimulates glucagon secretion remains unknown. Here, we show that arginine-vasopressin (AVP), secreted from the posterior pituitary, stimulates glucagon secretion. Glucagon-secreting alpha-cells express high levels of the vasopressin 1b receptor gene (*Avpr1b*). Activation of AVP neurons *in vivo* increased circulating copeptin (the C-terminal segment of the AVP precursor peptide, a stable surrogate marker of AVP) and increased blood glucose; effects blocked by pharmacological antagonism of either the glucagon receptor or vasopressin 1b receptor. AVP also mediates the stimulatory effects of hypoglycemia produced by exogenous insulin and 2-deoxy-D-glucose on glucagon secretion. We show that the A1/C1 neurons of the medulla oblongata drive AVP neuron activation in response to insulin-induced hypoglycemia. Exogenous injection of AVP *in vivo* increased cytoplasmic Ca^2+^ in alpha-cells (implanted into the anterior chamber of the eye) and glucagon release. Hypoglycemia also increases circulating levels of AVP in humans and this hormone stimulates glucagon secretion from isolated human islets. In patients with T1D, hypoglycemia failed to increase both plasma copeptin and glucagon levels. These findings suggest that AVP is a physiological systemic regulator of glucagon secretion and that this mechanism becomes impaired in T1D.

## Introduction

Glucagon is secreted from alpha-cells of the pancreatic islets and has many physiological actions; most notably the potent stimulation of hepatic glucose production to restore euglycemia when blood glucose has fallen below the normal range (a process referred to as counter-regulation). The importance of glucagon for glucose homeostasis is well established (1). In both type-1 diabetes (T1D) and type-2 diabetes (T2D), hyperglycemia results from a combination of complete/partial loss of insulin secretion and over-secretion of glucagon (2). In addition, counter-regulatory glucagon secretion becomes impaired in both forms of diabetes (particularly T1D), which may result in life-threatening hypoglycemia (3). Despite the centrality of glucagon to diabetes etiology, there remains considerable uncertainty about the regulation of its release and the relative importance of intra-islet effects and systemic factors (4, 5). Based on observations in isolated (*ex vivo*) islets, hypoglycemia has been postulated to stimulate glucagon secretion via intrinsic (6–8) and/or paracrine mechanisms (9, 10). While it is indisputable that the islet is a critical component of the body’s ‘glucostat’ (11) and has the ability to intrinsically modulate glucagon output, it is clear that such an ‘islet-centric’ viewpoint is overly simplistic (12). Indeed, many studies have clearly demonstrated that brain-directed interventions can profoundly alter islet alpha-cell function, with glucose-sensing neurons in the hypothalamus being key mediators (13–16). This ability of the brain to modulate glucagon secretion is commonly attributed to autonomic innervation of the pancreas (14, 17, 18). However, glucagon secretion is not only restored in pancreas transplantation patients but also insensitive to adrenergic blockade (19, 20), suggesting that other (non-autonomic) central mechanisms may also regulate glucagon secretion *in vivo*.

Arginine-vasopressin (AVP) is a hormone synthesised in the hypothalamus (reviewed in (21)). AVP neurons are divided into two classes: parvocellular AVP neurons (which either project to the median eminence to stimulate ACTH and glucocorticoid release or project centrally to other brain regions) and magnocellular AVP neurons (which are the main contributors to circulating levels of AVP). The parvocellular neurons reside solely in the paraventricular nucleus of the hypothalamus (PVH), whereas magnocellular neurons are found in both the PVH and supraoptic nucleus (SON). Stimulation of the magnocellular AVP neurons causes release of AVP into the posterior pituitary, where it enters the peripheral circulation.

Under normal conditions, *Avpr1b* is one of the most enriched transcripts in alpha-cells from both mice and humans (22, 23). This raises the possibility that AVP may be an important regulator of glucagon secretion under physiological conditions. Indeed, the ability of exogenous AVP and AVP analogues to potently stimulate glucagon secretion *ex vivo* has been known for some time (24). However, whether circulating AVP contributes to physiological counter-regulatory glucagon release and how this regulation is affected in diabetes remains unknown.

Here, we have investigated the regulation of glucagon by circulating AVP. We first explored the role of AVP during hypoglycemia, a potent stimulus of glucagon secretion. Next, we explored the link between AVP and glucagon in humans and provide evidence that this putative ‘hypothalamic-alpha-cell axis’ is impaired in T1D.

## Results

### AVP evokes hyperglycemia and hyperglucagonemia

We first investigated the metabolic effects of AVP *in vivo* (**Figure 1a-c**). We expressed the modified human M3 muscarinic receptor hM3Dq (see (25)) in AVP neurons by bilaterally injecting a Cre-dependent virus containing hM3Dq (AAV-DIO-hM3Dq-mCherry) into the supraoptic nucleus (SON) of mice bearing *Avp* promoter-driven Cre recombinase (*Avp*^ires-Cre+^ mice; **Figure 1a**). Expression of hM3Dq was limited to the SON (**Supplementary Figure 1a-c**), thus allowing targeted activation of magnocellular AVP neurons (that release AVP into the circulation). Patch-clamp recordings confirmed that bath application of clozapine-N-oxide (CNO; 5-10 µM) – a specific, pharmacologically inert agonist for hM3Dq – induced membrane depolarisation and increased the firing rate in hM3Dq-expressing AVP neurons (**Supplementary Figure 1d,e**). Injection of CNO (3 mg/kg i.p.) *in vivo* increased blood glucose (**Figure 1b**). We measured copeptin, the C-terminal segment of the AVP precursor peptide, which is a stable surrogate marker for AVP (26, 27), but the sample volume requirements (100 µL plasma) only allowed a single (terminal) measurement. Despite these experimental limitations, copeptin was elevated in response to CNO compared to saline injection (**Figure 1c**). It is notable that copeptin is a much larger peptide than AVP and its clearance from the circulation is slower (28). Thus, circulating AVP levels may undergo more dramatic variations (see (29) and data shown below).

**Figure 1:**
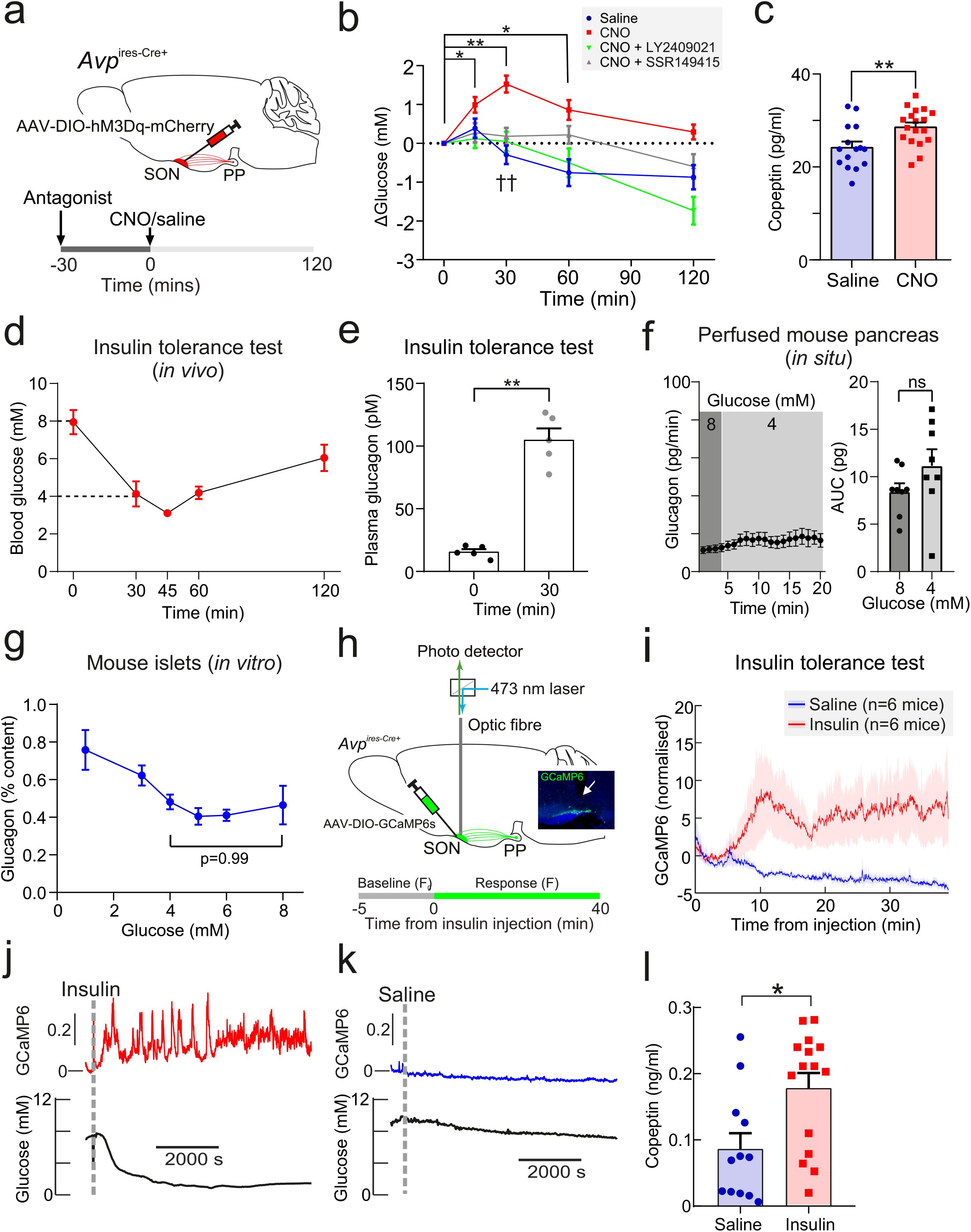
Insulin-induced hypoglycemia enhances population activity of AVP neurons in the SON, driving glucagon secretion. (a) AAV-DIO-hM3Dq-mCherry was injected bilaterally into the supraoptic nucleus (SON) of *Avp*^ires-Cre+^ mice. CNO (3 mg/kg) or vehicle was injected i.p. In the same cohort (different trial), LY2409021 (5 mg/kg) or SSR149415 (30 mg/kg) was injected (i.p.) 30 minutes prior to CNO. See **Supplementary Figure 1**. (b) Blood glucose measurements from (a). Two-way RM ANOVA (Tukey’s). CNO 0 mins vs. CNO at 15, 30 and 60 mins; p<0.05=*, p<0.01=**. Comparison of CNO vs. Saline, CNO+LY2409021 or CNO+SSR149415 at 30 mins; p<0.01=††. Time, p<0.0001; Treatment, p=0.0006; Interaction, p<0.0001. n=6 mice. (c) Terminal plasma copeptin 30 mins following saline or CNO injection. Mann Whitney t-test (p=0.0025, **). n=15-18 mice. (d) Plasma glucose during an insulin tolerance test (ITT; 0.75 U/kg) in n=5 wild-type mice. (e) Plasma glucagon following an ITT. n=5 wild-type mice. Paired t-test, p<0.01=**. (f) Glucagon secretion from the perfused mouse pancreas. *Right*: area under curve. Paired t-test, ns=not significant. (g) Glucagon secretion from isolated mouse islets. n=7 wild-type mice. One-way ANOVA with Tukey post-hoc. 4 mM vs 8 mM glucose, p=0.99. (h) Measurements of population GCaMP6s activity in pituitary-projecting AVP neurons in the supraoptic nucleus (SON). *Inset:* Expression of GCaMP6s in AVP neurons in the SON. Arrow = tip of the optic fiber. (i) GCaMP6s signal (normalized) in response to insulin (n=6 mice) or saline vehicle (n=6). (j) Simultaneous *in vivo* fiber photometry of AVP neuron activity (GCaMP6) and continuous glucose monitoring (black line) in response to an ITT (1 U/kg). Dashed grey line indicates the time of insulin injection. (k) Same animal as in (j), but for saline vehicle injection (dashed grey line). (l) Plasma copeptin at 30 mins following saline or insulin. Mann-Whitney U test, p=0.021. n=15-18 mice.

To establish the contribution of glucagon to this hyperglycemic response of SON AVP neuron activation, we pre-treated mice with the glucagon receptor antagonist LY2409021 (30). This completely abolished the hyperglycemic action of CNO (**Figure 1b**). Similarly, to understand the contribution of vasopressin 1b receptor (V1bR) signalling, we pre-treated mice with the V1bR antagonist SSR149415 (31). This also abolished the hyperglycemic effect of CNO (**Figure 1b**), suggesting that V1bR signalling mediates this response. CNO did not change food intake (**Supplementary Figure 1f**) and did not have an off-target effect on blood glucose in *Avp*^ires*-*Cre+^ mice expressing a passive protein (mCherry) under AAV transfection in the SON (**Supplementary Figure 1g**). Exogenous AVP also caused an increase in glucose (measured by continuous glucose monitoring or standard blood sampling) and glucagon relative to saline injection in wild-type mice (**Supplementary Figure 1h-j**).

### Insulin-induced glucagon secretion *in vivo* is driven by AVP

We next investigated whether AVP stimulates glucagon secretion during hypoglycemia *in vivo*. Insulin-induced hypoglycemia (from 8 to 4 mM) increased circulating glucagon levels 10-fold (**Figure 1d,e**). The same decrease in extracellular glucose only marginally stimulated glucagon secretion measured in the perfused mouse pancreas (**Figure 1f**). Similarly, reducing the glucose concentration from 8 to 4 mM does not stimulate glucagon secretion from isolated (*ex vivo*) islets (**Figure 1g**). This glucose dependence of glucagon secretion in isolated islets and the perfused pancreas is in keeping with numerous other reports (5, 32–36). Therefore, additional mechanisms extrinsic to the islet clearly participate in the control glucagon secretion *in vivo*.

We hypothesised that this extrinsic stimulus involves AVP. We investigated (using *in vivo* fiber photometry) whether AVP neuron activity is increased in response to an ITT (**Figure 1h**). Hypoglycemia (induced by insulin) evoked an increase in AVP neuron activity, whereas saline vehicle treatment was ineffective (**Figure 1i**). To investigate the glucose-dependence of the AVP neuron response to insulin, we simultaneously recorded AVP neuron activity and plasma glucose (by continuous glucose monitoring; **Figure 1j,k**). This revealed that the initial peak in AVP neuron activity following insulin injection occurs when blood glucose has fallen to 4.9±0.4 mM glucose (**Figure 1j** and **Supplementary Figure 2**). We measured plasma copeptin in response to insulin-induced hypoglycemia. Again, these experiments were terminal due to copeptin sample volume requirements. In these experiments, copeptin was increased by 114% (**Figure 1l**).

### AVP stimulates glucagon-secreting pancreatic alpha-cells

To understand how AVP increases glucagon secretion, we characterised its effect on isolated islets. Mouse islets express mRNA for the vasopressin 1b receptor (V1bR; encoded by *Avpr1b*), whereas vasopressin receptor subtypes 1a and 2 mRNA (*Avpr1a* and *Avpr2*) were found in the heart and the kidneys, consistent with their distinct roles in the regulation of blood pressure and diuresis (21), respectively (**Supplementary Figure 3a**). To confirm that *Avpr1b* expression was enriched in alpha-cells, mice bearing a proglucagon promoter-driven Cre-recombinase (*Gcg*^Cre+^ mice) were crossed with mice expressing a Cre-driven fluorescent reporter (RFP). qPCR of the fluorescence-activated cell sorted RFP^+^ and RFP^−^ fractions revealed that expression of *Avpr1b* is high in alpha-cells (RFP+) with ∼43-fold enrichment above that seen in RFP-cells (principally beta-cells) (**Supplementary Figure 3b,c**).

We explored whether the hyperglycemic and hyperglucagonemic actions of AVP are due to AVP directly stimulating glucagon secretion from alpha-cells. In dynamic measurements using the *in situ* perfused mouse pancreas, AVP produced a biphasic stimulation of glucagon secretion (**Figure 2a**). Glucagon secretion is a Ca^2+^-dependent process (37). We therefore crossed *Gcg*^Cre+^ mice with a Cre-dependent GCaMP3 reporter mouse (from hereon, *Gcg*-GCaMP3 mice), and implanted islets from these mice in the anterior chamber of the eye of recipient wild-type mice (**Figure 2b-d**; see (38)). This allowed the cytoplasmic Ca^2+^ concentration ([Ca^2+^]_i_) in individual alpha-cells to be imaged *in vivo*. Administration (i.v.) of AVP resulted in a biphasic elevation of [Ca^2+^]_i_ consisting of an initial spike followed by rapid oscillatory activity (**Figure 2c,d**), similar to the biphasic stimulation of glucagon secretion seen in the perfused pancreas.

**Figure 2:**
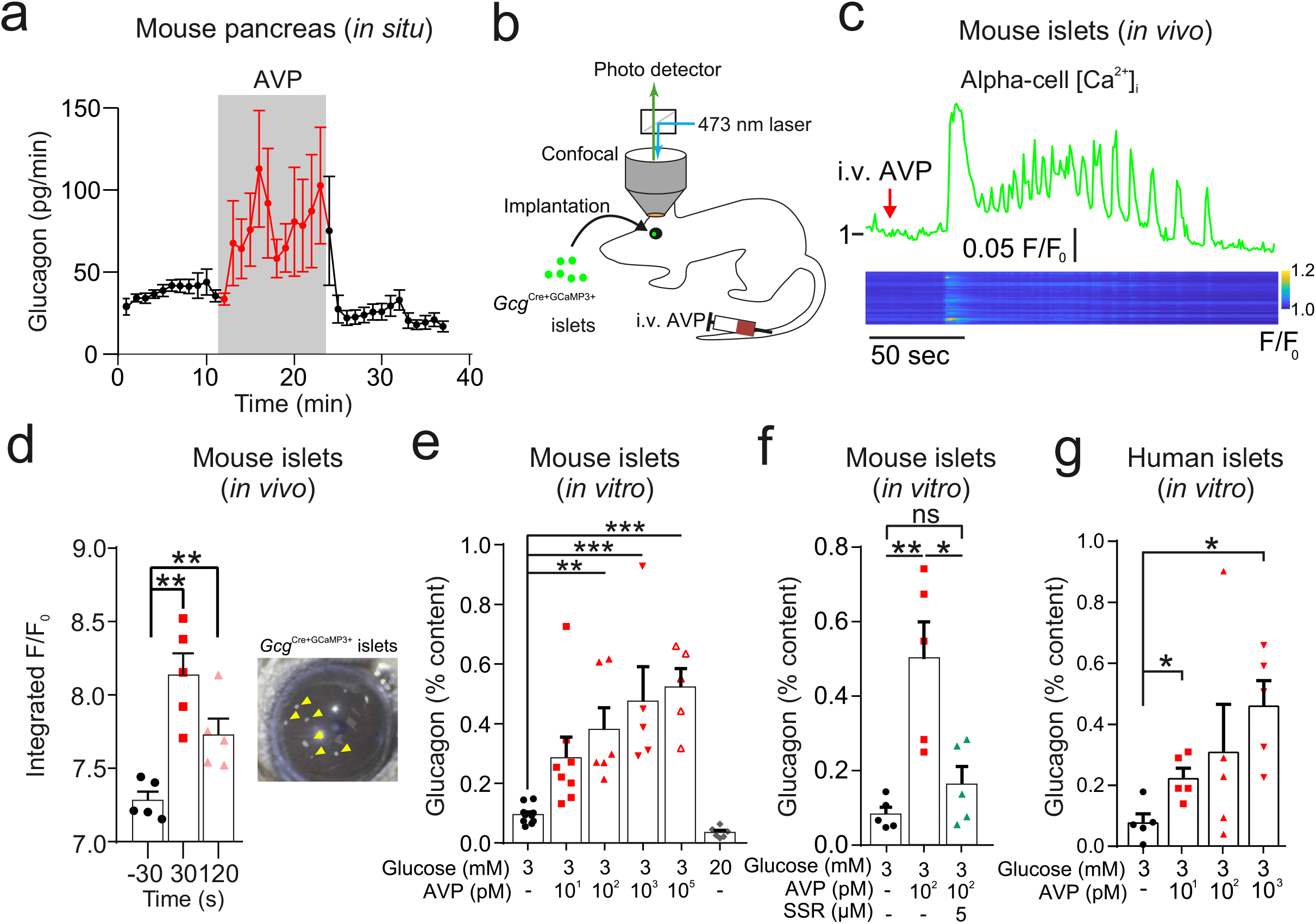
AVP increases glucagon release and alpha-cell activity *ex vivo*, *in situ* and *in vivo*. (a) Glucagon secretion from the perfused mouse pancreas in response AVP (10 nM). All data are represented as mean ± SEM. n=5 mice. Extracellular glucose of 3 mM. (b) Islets from *Gcg*^Cre+^-GCaMP3 mice were injected into the anterior chamber of the eye (ACE) of recipient mice (n=5 islets in 5 mice). After >4 weeks, GCaMP3 was imaged *in vivo* in response to i.v. AVP (10 µg/kg) or saline administration. Saline did not change the GCaMP3 signal. Signal is GCaMP3 fluorescence (F) divided by baseline signal (F_0_). AVP evoked an increase in calcium activity, typically starting with a large transient. *Below:* Raster plot of response (normalized F/F_0_) in different cells (ROIs) with a single islet. (c) Response of alpha-cell to i.v. AVP. Lower panel shows raster plot of response in different cells. (d) Integrated F/F_0_ (area under curve) response for all alpha-cells in recorded islets (5 islets, N=3 mice). The area under the curve was calculated 30 secs before i.v. injection, 30 secs after and 120 secs after. One-way RM ANOVA with Tukey’s multiple comparison test; p<0.01=**. *Right:* Image of islets (arrows) engrafted in the ACE. (e) Glucagon secretion from isolated mouse islets in response to AVP. One-way ANOVA (p<0.05=*; p<0.01=**; p<0.001=***). n=5-10 wild-type mice in each condition. (f) Glucagon secretion from isolated mouse islets in response to AVP in the presence and absence of the V1bR antagonist SSR149415. One-way ANOVA (p<0.05=*; p<0.01=**). n=5 wild-type mice per condition. (g) Glucagon secretion from islets isolated from human donors, in response to AVP. Paired t-tests, p<0.05=*. n=5 human donors.

In isolated mouse islets, AVP stimulated glucagon secretion (EC_50_ = 25.0 pM; 95% CI = (4.80, 133) pM) in islets incubated in 3 mM glucose (**Figure 2e**). The AVP-induced increase in glucagon was prevented by SSR149415 (**Figure 2f**). AVP also reversed the glucagonostatic effect of 15 mM glucose but higher concentrations (>1 nM) were required (**Supplementary Figure 4a**). AVP at concentrations up to 100 nM did not affect insulin secretion from mouse islets when tested at 3 mM or 15 mM glucose (**Supplementary Figure 4b,c**). Finally, AVP also stimulated glucagon secretion (EC_50_ = 7.69 pM; 95% CI = (5.10, 113) pM) in isolated human pancreatic islets (**Figure 2g**).

To understand the intracellular mechanisms by which AVP stimulates glucagon secretion, we isolated islets from *Gcg*-GCaMP3 mice. We first performed perforated patch-clamp recordings of membrane potential in intact islets from *Gcg*-GCaMP3 mice. AVP increased action potential firing frequency (**Figure 3a,b**). Next, we conducted confocal imaging of [Ca^2+^]_i_ in these islets and confirmed that AVP increased [Ca^2+^]_i_ in alpha-cells (**Figure 3c,d**). The capacity of AVP to increase [Ca^2+^]_i_ in alpha-cells was abolished following application of the V1bR antagonist SSR149415 (**Figure 3e**). AVP-induced Ca^2+^ activity was dependent on G_q_-protein activation, because it was blocked with the G_q_-inhibitor YM254890 ((39); **Figure 3f**). Furthermore, AVP increased intracellular diacylglycerol, which is a downstream product of G_q_ activation (**Figure 3g,h**).

**Figure 3:**
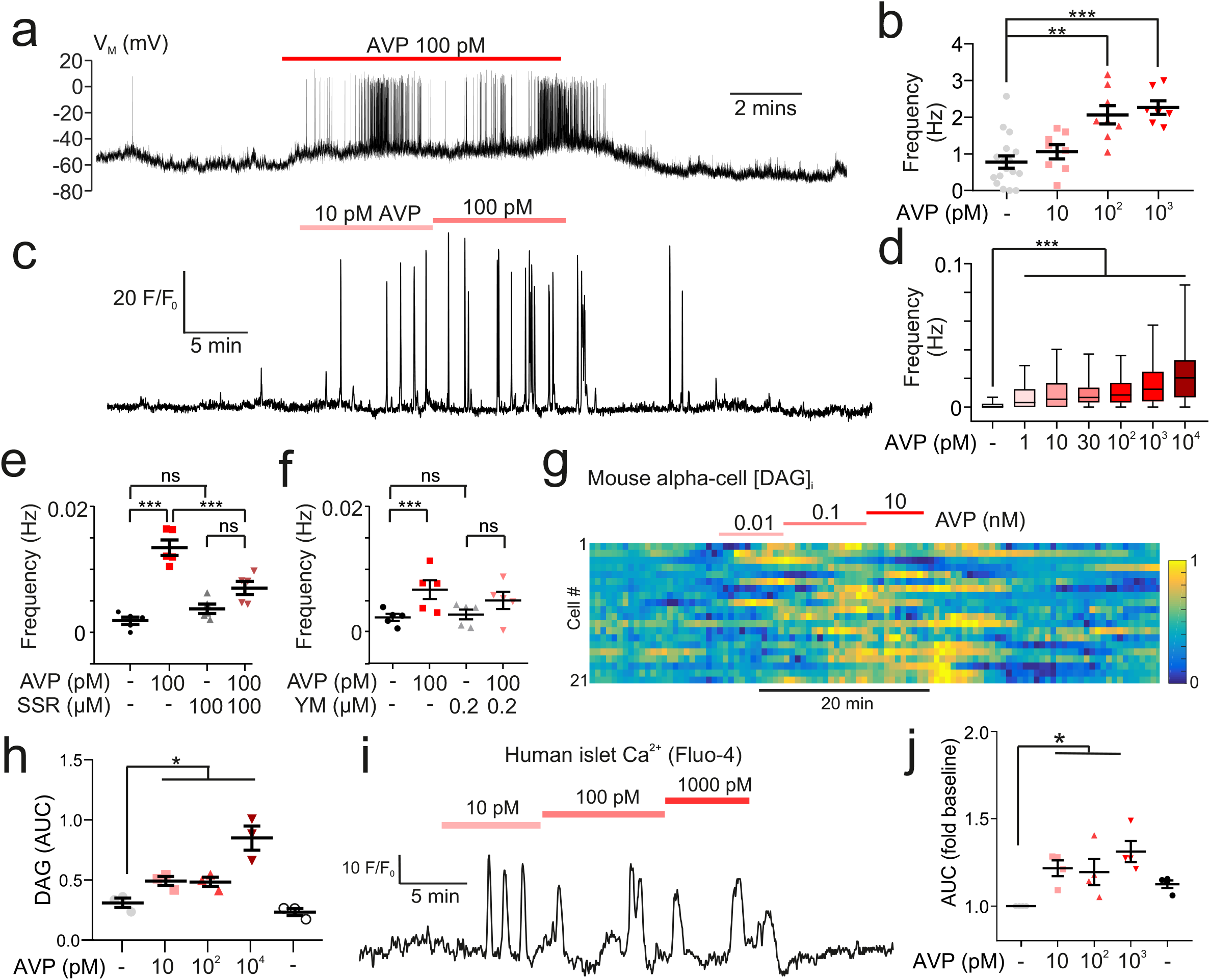
AVP increases action potential firing, Ca^2+^ activity and intracellular DAG in alpha-cells in intact islets. (a) Membrane potential (V_m_) recording (perforated patch-clamp) of an alpha-cell in response to 100 pM AVP. (b) Frequency-response curve for varying concentrations of AVP (17 alpha-cells, 10 *Gcg*-GCaMP3 mice). Mixed-effects analysis of variance, Holm-Sidak’s post-hoc (p<0.01=**; p<0.001=***; p=0.073 for 3 mM glucose vs. 10 pM AVP). (c) GCaMP3 signal from an alpha-cell in response to AVP. (d) Box and whisker plot of the frequency of GCaMP3 oscillations in response to AVP. 142-170 alpha-cells, 7 islets, n=7 *Gcg-*GCaMP3 mice. Recordings in 3 mM glucose. One-way RM ANOVA, p<0.001=***. (e) Frequency of GCaMP3 oscillations in response to 100 pM AVP in the presence and absence of SSR149415 (10 µM). 75-90 alpha-cells, 6 islets, n=5 *Gcg*-GCaMP3 mice. Recordings in 3 mM glucose. One-way ANOVA (Tukey), p<0.001=***, ns=not significant (p>0.2). (f) Frequency of GCaMP3 oscillations in response to 100 pM AVP in the absence and presence of YM-254890 (0.2 µM). 75-90 alpha-cells, 6 islets, n=5 *Gcg*^Cre+^-GCaMP3 mice. Recordings in 3 mM glucose. One-way RM ANOVA (Tukey’s multiple comparisons test), p<0.05=*, ns=not significant (p>0.3). (g) Heatmap of intracellular diacylglycerol (DAG; Upward DAG) signal from single islet cells (dispersed into clusters) in response to AVP. The signal was median filtered and normalized to largest signal in the recording. (h) Area under curve (AUC, normalized to duration) for DAG signal for each experimental condition. 10 recordings, 152 cells, n=3 wild-type mice. One-way RM ANOVA, p<0.05=* (Tukey’s multiple comparisons test). (i) Fluo4 signal from a putative alpha-cell in a human islet in response to AVP (10, 100 and 1000 pM). Recording in 3 mM glucose. (j) Area under curve (AUC, normalized to duration) for Fluo4 signal in each human islet, for each experimental condition. 26 islets, n=4 human donors. One-way ANOVA, p<0.05=* (Sidak).

In human islets, *AVPR1B* was the most abundant of the vasopressin receptor family (**Supplementary Figure 3d**). These data are supported both by a recent meta-analysis of single-cell RNA-seq data from human donors (40), and bulk sequencing of human islet fractions (41). Finally, in alpha-cells in human islets (identified by their response to adrenaline (42)), AVP increased [Ca^2+^]_i_ (**Figure 3i,j**).

### Pharmacological and genetic inhibition of AVP signaling suppress counter-regulatory glucagon secretion

Given the relatively small increase in circulating copeptin in response to hypoglycemia (**Figure 1l**), we used pharmacological and genetic approaches to more conclusively establish the link between AVP and counter-regulatory glucagon during hypoglycemia. We injected wild-type mice with the V1bR antagonist SSR149415 prior to an insulin tolerance test (ITT) (**Figure 4a,b**). This reduced glucagon secretion during insulin-induced hypoglycemia by 60%. Similarly, in *Avpr1b* knockout mice (*Avpr1b*^−/−^; (43)) glucagon secretion was decreased by 65% compared to wild-type littermates (*Avpr1b*^−/−^; **Figure 4c,d**). Despite the drastic reduction in plasma glucagon by pharmacological antagonism of the V1bR or genetic KO of *Avpr1b*, the depth of hypoglycemia was not affected (**Figure 4a,c**). Insulin measured during an ITT revealed that circulating insulin increases from a basal of 110±12 pM to 910±74 pM at 30 min (n=5 mice). Thus, insulin is present at an ∼10-fold molar excess compared to glucagon. This explains why the hypoglycemic effect of exogenous insulin predominates in this experimental paradigm, with no change in plasma glucose despite a strong reduction in circulating glucagon. Indeed, in mice an ITT tests both counter-regulation and insulin sensitivity as recently reviewed (44).

**Figure 4:**
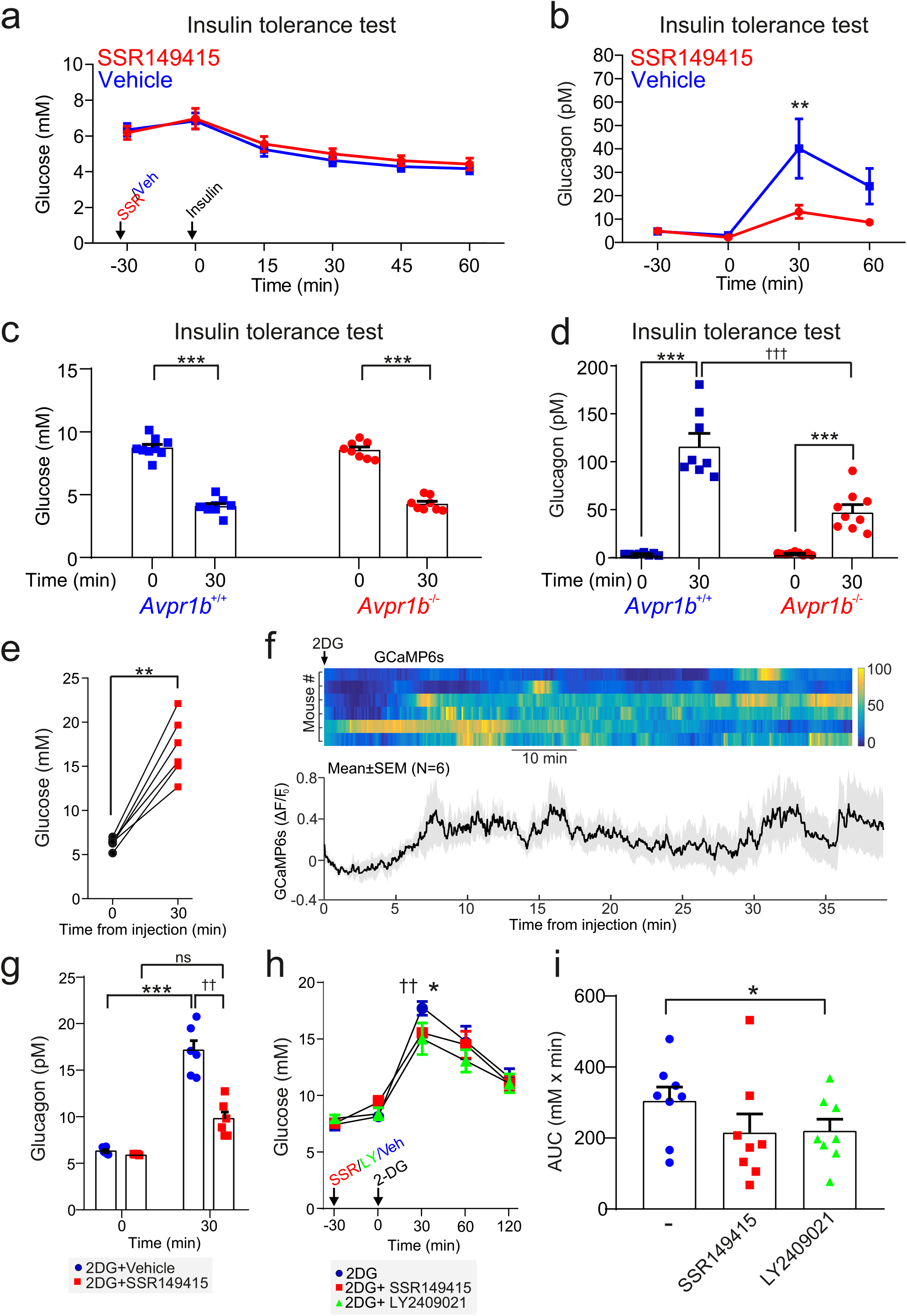
The vasopressin 1b receptor mediates hypoglycemia-induced glucagon secretion. (a) Blood glucose during an ITT (0.75 U/kg; injection at 30 min). 30 mins prior to the commencement of the ITT (t=0 min), either the V1bR antagonist SSR149415 (30 mg/kg) or vehicle was administered i.p. n=7 wild-type mice. (b) Plasma glucagon for (a). Two-way RM ANOVA with Sidak’s (between conditions) and Tukey’s (within condition) multiple comparisons test. Vehicle vs. SSR149415; p<0.01=†† (30 minutes) and p<0.05=† (60 minutes). Glucagon was increased in response to an ITT in both treatment groups (p<0.01=** vs. 0 min). n=6-7 wild-type mice. (c) Plasma glucose during an ITT (0.75 U/kg; injection at 0 min) in *Avpr1b*^−/−^ mice and littermate controls (*Avpr1b*^+/+^). Two-way RM ANOVA (Tukey), p<0.001=***. n=8-9 mice. (d) Plasma glucagon for (c). Two-way RM ANOVA (Sidak). *Avpr1b*^−/−^ vs. *Avpr1b*^+/+^; p<0.001=††† (30 minutes). 0 vs. 30 min; p<0.001=***. n=8-9 mice. (e) Population GCaMP6s activity in pituitary-projecting AVP neurons in the supraoptic nucleus (SON). GCaMP6s was imaged in response to 2-Deoxy-D-glucose (2DG, 500 mg/kg) injection (i.p.). Plasma glucose at baseline (0 min) and 30 min after 2DG injection. Paired t-test, p<0.01=**. n=6 mice. (f) *Upper:* heatmap of population activity (GCaMP6s) response to 2DG for each mouse (n=6). *Lower:* mean ± SEM GCaMP6s signal for all mice (n=6) in response to 2DG. GCaMP6s data represented as (F-F_0_)/F_0_. (g) Plasma glucagon following 2DG injection (at 0 min). Prior to 2DG, either SSR149415 or vehicle was administered i.p. Two-way RM ANOVA by both factors (Bonferroni). Vehicle vs. SSR149415; p<0.01=†† (30 mins); p>0.99 (0 mins). 0 vs. 30 mins; p=0.009 (Vehicle); p=0.093 (SSR149415). Time, p<0.0001; Treatment, p=0.002; Interaction, p=0.005. n=6 mice. (h) Blood glucose response to 2DG, with or without pre-treatment with SSR149415 (30 mg/kg), LY2409021 (5 mg/kg) or saline vehicle. Antagonists/vehicle injected 30 mins prior to 2DG. Two-way RM ANOVA with Sidak’s multiple comparison test; 2DG vs. 2DG+SSR149415, p=0.0103 (*); 2DG vs. 2DG+LY024091, p=0.0047 (††). (i) Area under the (glucose) curve (AUC). One way ANOVA; 2DG vs. 2DG+LY024091, p<0.05 (*).

We also tested the ability of AVP to modulate glucagon *in vivo* in response to 2-deoxy-D-glucose (2DG). 2DG is a non-metabolizable glucose molecule that evokes a state of perceived glucose deficit (mimicking hypoglycemia) and triggers a robust counter-regulatory stimulation of glucagon secretion (17). We monitored AVP neuron activity in response to 2DG by *in vivo* fiber photometry, and correlated this to changes in plasma glucose during this metabolic challenge. Injection (i.p.) of 2DG increased blood glucose (**Figure 4e**) and triggered a concomitant elevation of [Ca^2+^]_i_ in AVP neurons (**Figure 4e-h**). The elevation in plasma glucagon by 2DG injection was attenuated by 50% following pretreatment with the V1bR antagonist SSR149415 (**Figure 4g**). The hyperglycemic response to 2DG was also partially antagonized by pre-treatment with either the V1bR antagonist SSR149415 or the glucagon receptor antagonist LY2409021 (**Figure 4h,i**). We conclude that AVP contributes to the hyperglycemic response to 2DG, and it does so (at least in part) by stimulating glucagon release.

### Hypoglycemia evokes AVP secretion via activation of A1/C1 neurons

Many physiological stressors activate hindbrain catecholamine neurons, which release noradrenaline (A1) or adrenaline (C1) and reside in the ventrolateral portion of the medulla oblongata (VLM). Activation of C1 neurons (by targeted glucoprivation or chemogenetic manipulation) elevates blood glucose (45–47) and plasma glucagon (48). Furthermore, C1 cell lesions severely attenuate the release of AVP in response to hydralazine-evoked hypotension (49), indicating that this hindbrain site may be a key regulator of AVP neuron activity during physiological stress.

To characterize any functional connectivity between A1/C1 neurons and SON AVP neurons, we conducted channelrhodpsin-2-assisted circuit mapping (CRACM; (50)) and viral tracer studies (**Figure 5a-e**). We injected a Cre-dependent viral vector containing the light-gated ion channel channelrhodopsin-2 (AAV-DIO-ChR2-mCherry) into the VLM (targeting A1/C1 neurons) of *Avp^GFP^ x Th*^Cre+^ mice (**Figure 5a**). Projections from the A1/C1 neurons were present in the SON and PVH and co-localised with AVP-immunoreactive neurons (**Figure 5b** and **Supplementary Figure 5a**). A1/C1 neurons express vesicular glutamate transporter 2 and as a result co-release glutamate with catecholamines (see review (51)). Therefore, by monitoring glutamatergic excitatory post-synaptic currents (EPSCs) evoked by ChR2 activation, we could determine whether A1/C1 neurons are synaptically connected to AVP neurons. Brain slice electrophysiology revealed that in the majority (89%) of GFP^+^ AVP neurons, opto-activation of A1/C1 neuron terminals results in EPSCs (**Figure 5a-e**). These EPSCs were glutamatergic, as they were abolished by the AMPA and kainate receptor antagonist DNQX (**Figure 5c**). Furthermore, these EPSCs could be blocked with TTX, but reinstated with addition of 4-AP (**Figure 5e**), indicating that A1/C1 neurons connect to AVP neurons in the SON monosynaptically.

**Figure 5:**
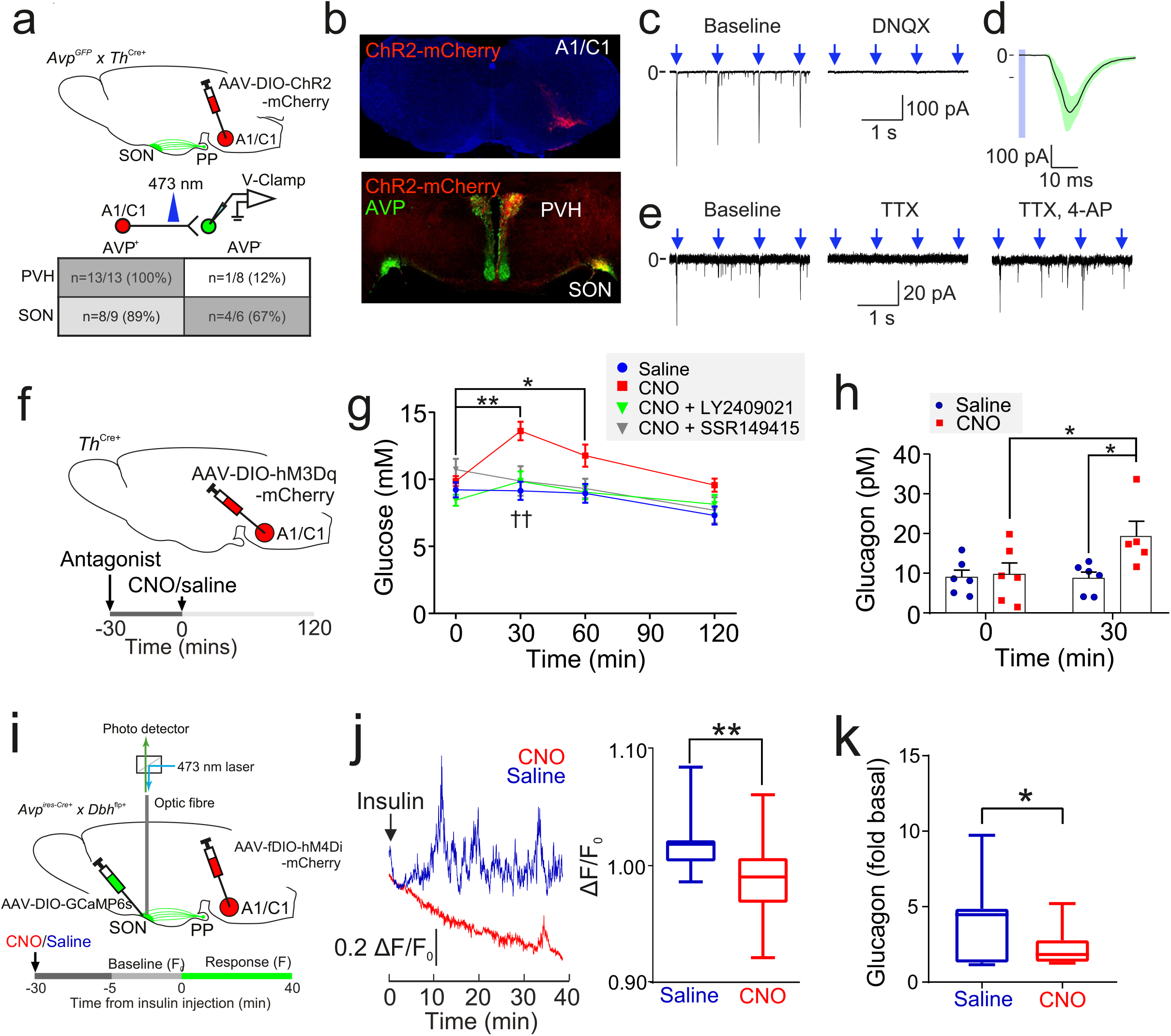
Insulin-induced AVP secretion is mediated by A1/C1 neurons. (a) *Upper:* AAV-DIO-ChR2-mCherry was injected into the VLM of *Th*^Cre+^ mice or *Avp*^GFP^ x *Th*^Cre+^ mice, targeting A1/C1 neurons. *Lower:* CRACM. Excitatory post-synaptic currents (EPSCs) were recorded in voltage-clamp mode in GFP^+^ (AVP) and GFP^−^ neurons in the SON and PVH. The number (n) of neurons that responded to opto-activating A1/C1 terminals. Total of 36 neurons recorded from 5 mice. See **Supplementary Figure 5a**. (b) *Upper:* Viral expression of ChR2-mCherry in A1/C1 neurons. *Lower:* A1/C1 neuron terminals co-localise with AVP-immunoreactive neurons. Representative from 3 mice. (c) *Left:* EPSCs evoked by opto-activation of A1/C1 terminals with 473 nm light pulses (arrows). *Right:* Light-evoked EPSCs following application of DNQX (20 µM). Representative of 5 recordings from 3 mice. (d) EPSC waveforms in a single GFP^+^ (AVP) neuron in response to repeated opto-activation of A1/C1 neuron terminals. Black line and shaded are = Mean±SD of EPSCs. Light pulse = blue bar. Representative of 3 recordings from 3 mice. (e) EPSCs evoked by opto-activating A1/C1 terminals at baseline (*left*) and following addition of TTX (1 µM*; middle*) and 4-AP (1 mM*; right*). Representative of 3 recordings from 3 mice. (f) AAV-DIO-hM3Dq was injected into *Th*^Cre+^ mice, targeting A1/C1 neurons. CNO (1 mg/kg) was then injected (i.p.). Antagonists (or vehicle) for the V1bR (SSR149415, 30 mg/kg) or glucagon receptor (GCGR; LY2409021, 5 mg/kg) were injected 30 minutes prior to CNO. Plasma glucose and glucagon was then measured. See **Supplementary Figure 5b,c**. (g) Plasma glucose in response to CNO and pre-treatment with antagonists. n=8 mice. Two-way RM ANOVA (Sidak’s multiple comparison’s test). Time (p<0.0001), Treatment (p=0.03) and Interaction (p=0.0002).(h) Plasma glucagon at 30 mins post CNO (or vehicle) injection. Two-way RM ANOVA with Tukey’s (within treatment) and Sidak’s (between treatments) multiple comparisons. p<0.05=*, ns=not significant. Within treatment, CNO increased glucagon at 30 min vs. 0 min (p=0.022). Saline did not (p=0.96). Between treatments, CNO increased glucagon at 30 mins vs. saline (p=0.001). n=6 mice. (i) *In vivo* fiber photometry measurements of population GCaMP6 activity in pituitary-projecting SON AVP neurons during A1/C1 neuron inhibition. AAV-DIO-GCaMP6s was injected into the SON and AAV-fDIO-hM4Di-mCherry into the VLM of *Avp*^ires-Cre+^ x *Dbh*^flp+^ mice. GCaMP6s was then imaged in response to an insulin tolerance test (ITT), following inhibition of the A1/C1 neuron (with CNO at 1 mg/kg), as indicated by the protocol in the lower horizontal bar. See **Supplementary Figure 6c**. (j) *Left:* Example population activity in one mouse (as described in (i)) in response to an ITT, following saline or CNO treatment (on different trials). CNO strongly inhibited the response to insulin. *Right:* Average GCaMP6 signal ((F-F_0_)/F_0_) during response to insulin with either saline or CNO pre-treatment (n=9 mice). CNO reduces the AVP GCaMP6 signal. t-test, p<0.01=**. (k) Plasma glucagon in response to an ITT in mice described in (i). 30 min before the insulin injection, either saline or CNO was given i.p. Glucagon is represented as fold of basal, where basal is 0 min (just prior to insulin) and the sample was taken at 30 mins post-insulin. t-test, p=0.023 (*). n=8 mice.

To explore the consequences of A1/C1 activation *in vivo*, we injected AAV-DIO-hM3Dq-mCherry bilaterally into the VLM of *Th*^Cre+^ mice (**Figure 5f** and **Supplementary Figure 5b,c**). Activation of A1/C1 neurons with CNO evoked a ∼4.5 mM increase in plasma glucose (**Figure 5g**). Pre-treatment with the glucagon receptor antagonist LY2409021 inhibited this response (**Figure 5g**). In line with this, plasma glucagon was increased following CNO application (**Figure 5h**). The hyperglycemic response was also dependent on functional V1bRs, because it was abolished following pre-treatment with the V1bR antagonist SSR149415 (**Figure 5g**). CNO had no effect on blood glucose in *Th*^Cre+^ mice expressing mCherry in A1/C1 neurons (**Supplementary Figure 5c**). Together, these data indicate that A1/C1 neuron activation evokes AVP release, which in turn stimulates glucagon secretion. We therefore hypothesised that hypoglycemia-induced AVP release is due to projections from A1/C1 neurons. In support of this hypothesis, c-Fos expression (a marker of neuronal activity) was increased in A1/C1 neurons following an insulin bolus (**Supplementary Figure 6a,b**).

To determine the contribution of the A1/C1 region to AVP neuron activity during an ITT, we inhibited the A1/C1 region whilst monitoring AVP neuron activity. To this end, we expressed an inhibitory receptor (the modified human muscarinic M4 receptor hM4Di; (52)) in A1/C1 neurons (by injecting AAV-fDIO-hM4Di-mCherry into the VLM) and GCaMP6s in AVP neurons (by injecting AAV-DIO-GCaMP6s into the SON) of *Dbh*^flp+^ x *Avp*^ires-Cre+^ mice (**Figure 5i-k** and **Supplementary Figure 6c**). We then measured AVP neuron population [Ca^2+^]_i_ activity (with *in vivo* fiber photometry) and plasma glucagon following inhibition of A1/C1 neurons with CNO. AVP neuron population activity during an ITT was partially reduced by A1/C1 silencing compared to vehicle injection (**Figure 5j**). Injection of CNO did not produce a statistically significant increase in basal plasma glucagon compared to saline (8±0.3 pM *vs.* 9±1 pM, respectively). However, when insulin was subsequently injected, glucagon rose significantly less in the CNO treated group (2-fold *vs.* 4-fold; **Figure 5k**). Together, these data suggest that A1/C1 neurons contribute to AVP-dependent glucagon secretion during an ITT.

### Insulin-induced AVP secretion underlies counter-regulatory glucagon secretion in humans

We extended these observations to human physiology. Healthy volunteers were given a hypoglycemic clamp during one visit and a euglycemic clamp during another visit in a randomized order (**Figure 6**). In response to hypoglycemia (blood glucose of 2.8±0.1 mM, **Figure 6a**), plasma glucagon increased by >300% (**Figure 6b**). In contrast, during euglycemia (blood glucose of 5.1±0.1 mM) glucagon was stable (**Figure 6b**). Measurements of plasma AVP revealed that AVP rose during a hypoglycemic clamp (from a basal 2 pM to 10 pM) but did not change during a euglycemic clamp (**Figure 6c**). There was a highly significant (p<0.001) correlation between AVP and glucagon (**Figure 6d**). Copeptin is more widely measured than AVP in clinical practice (53) and we therefore also measured the levels of this peptide. Like AVP, copeptin increased during the hypoglycemic clamp (**Figure 6e,f**). We compared copeptin and AVP measured in the same samples. Copeptin and AVP were significantly correlated but with a non-zero y-intercept (**Figure 6g**). This has been observed previously and is attributed to slower clearance kinetics of copeptin (28, 29). This is important as it suggests that using copeptin as a surrogate marker of AVP underestimates the true changes in AVP. Nevertheless, there was a linear and statistically significant correlation between glucagon and copeptin in the samples taken from both eu- and hypoglycemic clamps (**Figure 6h**).

**Figure 6:**
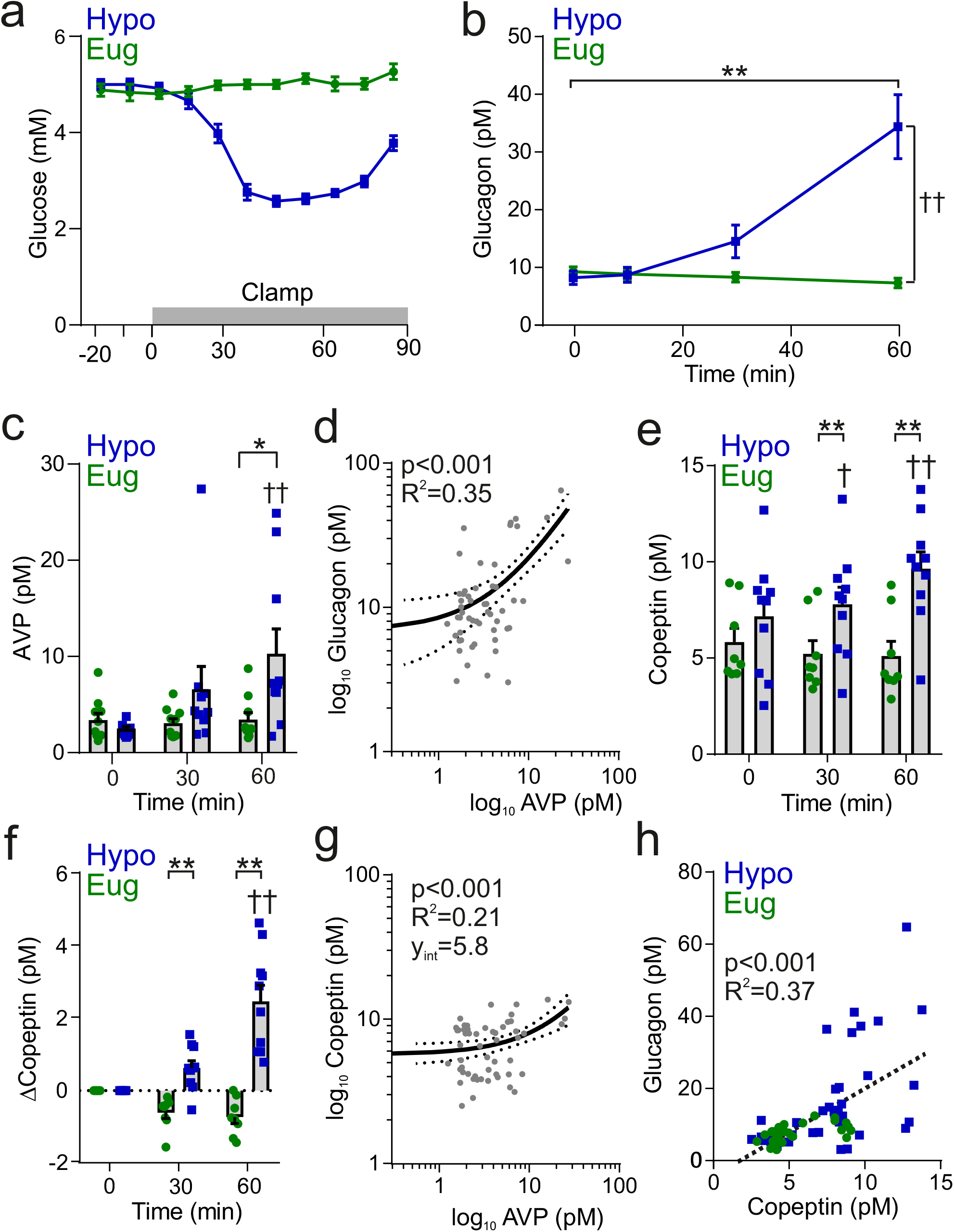
Insulin-induced hypoglycemia evokes copeptin and glucagon secretion in human participants. (a) Blood glucose was clamped at euglycemia (Eug) and followed during insulin-induced hypoglycemia (Hypo). n=10 healthy human subjects. The clamp was initiated at time 0 min and terminated at 60 min. (b) Plasma glucagon measurement during insulin-induced hypoglycemia. Two-way RM ANOVA by both factors. Time point *vs.* 0 min; p<0.01=**. Between treatments; p<0.01=††. (c) Plasma AVP measurement during and following clamping period. Two-way RM ANOVA by both factors. Hypoglycemia 0 min *vs* 60 min; p<0.01=††. Hypoglycemia 0 min *vs* 30 min; p=0.07. Between treatments; p<0.05=*. (d) Log-log plot of plasma AVP and plasma glucagon. Data points are 0, 30 and 60 min during hypoglycemic clamp. Linear regression (solid line) with 95% CI (dashed line). (e) Plasma copeptin measurement during hypoglycemic or euglycemic clamp. Two-way RM ANOVA by both factors. Indicated time point *vs.* 0 min; p<0.05=†, p<0.01=††. For hypoglycemic clamp 0 min vs. 30 min, p=0.07. Between treatments; p<0.01=**. (f) Change in plasma copeptin from baseline (time = 0 min). Two-way RM ANOVA by both factors. Indicated time point *vs.* 0 min; p<0.01=††. For hypoglycemic clamp 0 min vs. 30 min, p=0.07. Between treatments; p<0.01=**. (g) Log-log plot of plasma AVP and plasma copeptin (as plotted in (29)). Data points are 0, 30 and 60 min during hypoglycemic clamp. Linear regression (solid line) with 95% CI (dashed line). (h) Correlation of change in copeptin and glucagon, with a linear regression (dashed line). Data points are from both euglycemia and hypoglycemia at 0, 10, 30 and 60 min.

### Insulin-induced copeptin secretion is diminished in T1D

We measured copeptin and glucagon during hypoglycemic clamps in subjects with T1D and non-diabetic, BMI- and age-matched ‘control’ individuals (**Supplementary Table 1**). As expected, the T1D patients had higher plasma glucose levels than the healthy controls (**Figure 7a**). During the hypoglycemic clamp, blood glucose was reduced in both controls and T1D participants, with blood glucose converging at 2.8 mM after 60 minutes (**Figure 7a**). In control subjects, hypoglycemia evoked a ∼ 17 pM increase in plasma glucagon within 60 minutes (**Figure 7b**). In contrast, hypoglycemia failed to increase circulating glucagon in all subjects with T1D, even at 60 minutes (**Figure 7b**). For display, time-dependent changes in copeptin after induction of hypoglycemia are shown after subtraction of basal copeptin (7.0±0.8 pM and 4.5±0.6 pM in control and T1D participants, respectively). In line with the glucagon phenotype, copeptin was significantly elevated in controls but not in subjects with T1D (**Figure 7c**). We plotted plasma glucagon against total plasma copeptin. Overall, copeptin levels in the subjects with T1D clustered at the lower end of the relationship, and even when copeptin levels were similar between the groups, glucagon levels were lower in T1D subjects (**Figure 7d**).

**Figure 7:**
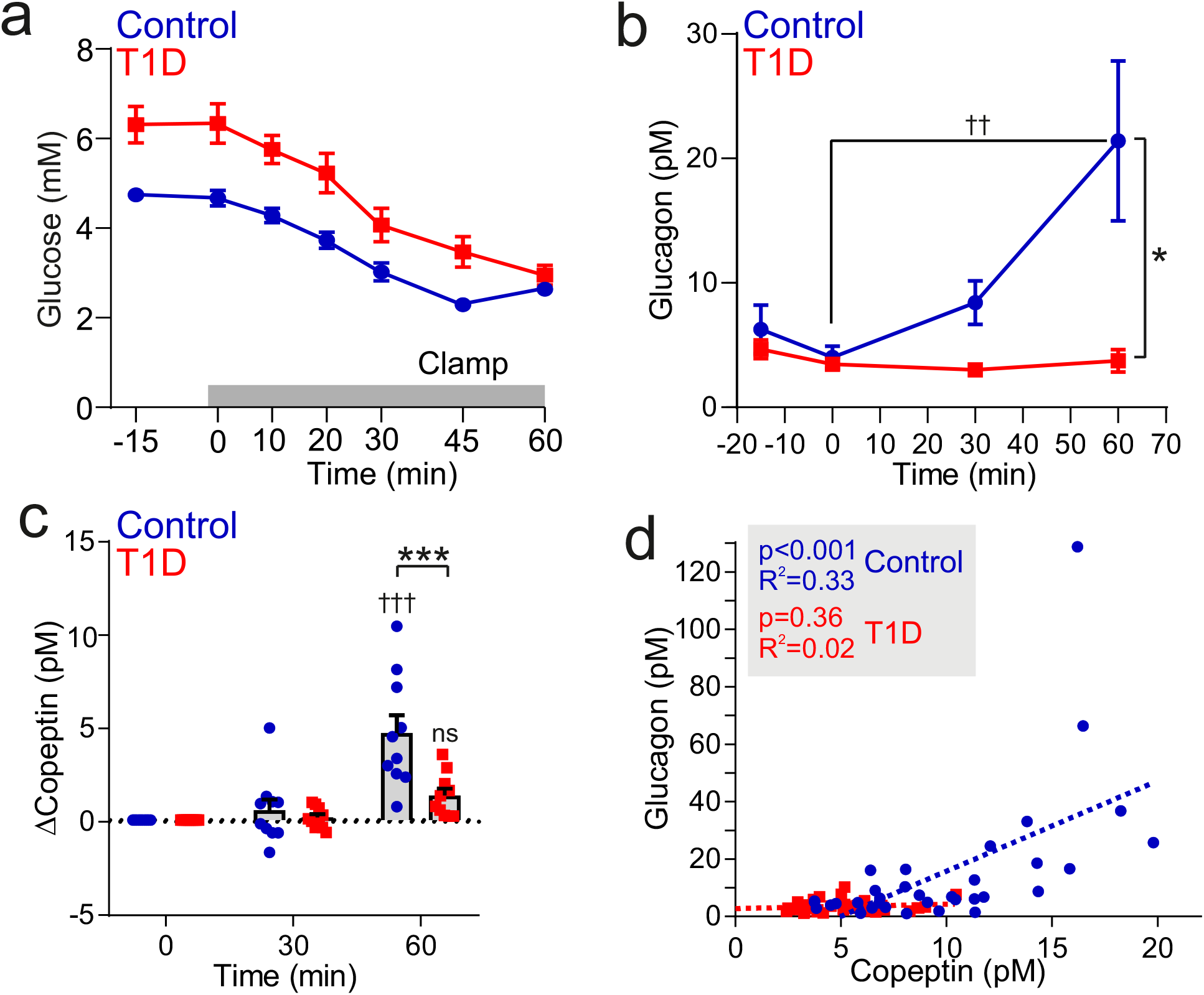
Insulin-induced copeptin and glucagon secretion is diminished in people with type 1 diabetes. (a) Hypoglycemia was induced by an insulin infusion in patients with T1D (n=10) and non-diabetic individuals (Controls; n=10). The insulin infusion was initiated at 0 min. (b) Plasma glucagon during the clamping period. Two-way RM ANOVA by both factors. Indicated time point *vs.* 0 min; p<0.01=††. Between groups; p<0.05=*. Data from n=10 Control and n=10 T1D. (c) Plasma copeptin measurement during and following clamping period. Change in copeptin from baseline (time = 0 min). Two-way RM ANOVA by both factors. Time point *vs.* 0 min; p<0.001=†††; ns = not significant (p>0.05). Between groups; p<0.001=***. Data from n=10 Control and n=10 T1D. (d) Correlation of copeptin and glucagon following hypoglycemic clamping for control participants (n=10, circle) and T1D (n=10, square). Linear regressions (dashed lines) for Control and T1D datasets.

## Discussion

We investigated the role of AVP in regulating glucagon secretion *in vivo* during hypoglycemia. In mouse, we first stimulated AVP neurons with a DREADD approach, and observed and increase in plasma glucose that was V1bR- and glucagon receptor-dependent. Administering AVP *in vivo* produced increases in circulating glucose and glucagon, and initiated [Ca^2+^]_i_ oscillations in alpha-cells transplanted into the anterior chamber of the eye. We also demonstrate that AVP neuron activity is elevated during hypoglycemia induced by either 2DG or exogenous insulin. Pharmacological antagonism or genetic KO of the V1bR revealed that AVP is a major contributor to counter-regulatory glucagon release.

We did not observe an effect of AVP on insulin secretion from isolated mouse islets, whether at low or high glucose. In contrast, earlier studies have demonstrated that AVP stimulates insulin secretion (24, 54–56). We attribute this discrepancy to the much higher (un-physiological) concentrations used in the earlier studies, which might have resulted in off-target effects. Indeed, 10 nM AVP directly closes ATP-sensitive K^+^ channels in insulin-secreting cell lines (57). It is notable that V1bR is the only receptor from the vasopressin receptor family expressed in mouse islets, and its expression is restricted to alpha-cells (58, 59), an observation we now confirm. It is possible that the previous reports of stimulatory effects of AVP on insulin secretion might reflect paracrine stimulation mediated by glucagon (60).

Catecholaminergic neurons in the VLM are a key component of the central counter-regulatory circuit and the ability of these neurons to evoke hyperglycemia (45–47) and hyperglucagonemia (48) is well-established. Recent studies have clearly demonstrated that spinally-projecting C1 neurons evoke hyperglycemia by stimulating the adrenal medulla (46, 47), suggesting that the response is likely to be mediated by corticosterone and/or adrenaline. We confirm that activation of A1/C1 neurons evokes hyperglycemia, but by promoting glucagon release. The important distinction here is that the elevation of plasma glucagon cannot be explained by adrenaline signalling, because circulating levels of adrenaline do not increase glucagon release (61). Neurons in the VLM have a well-documented ability to increase plasma AVP (49, 62). Our data strongly support the notion that the hyperglycemic and hyperglucagonemic effect of activating A1/C1 neurons is, at least in part, mediated by stimulation of AVP release: we show that A1/C1 neurons send functional projections to the SON, A1/C1 function is required for insulin-induced AVP neuron activation and that AVP is an important stimulus of glucagon secretion. However, we recognise that other circuits must be involved in the activation of AVP neurons, because A1/C1 neuron inhibition only partially prevented insulin-induced AVP neuron activation and glucagon release (see **Figure 5j,k**; although this may also be explained by variability in AAV-DIO-GCaMP6 and AAV-fDIO-hM4Di transfection in these experiments). For example, the PVN is richly supplied with axons from the BNST (63) and hypothalamic VMN neurons are key drivers of the glucose counter-regulatory response project to the BNST (15). Therefore, a VHN-BNST circuit may also be important for driving AVP neuron activity in response to hypoglycemia, explaining why A1/C1 inhibition only partially prevented the activation of AVP neurons. AVP infusion in human participants increases circulating glucagon (64), but it is also thought to directly stimulate glycogen breakdown from the liver (65). Consequently, it is possible that part of the increase in plasma glucose following A1/C1 neuron stimulation is mediated by the action of AVP on the liver.

The signal driving AVP neuron activation is unlikely to be due to a direct action of insulin on AVP or A1/C1 neurons, because 2-DG caused a similar activation. The widely accepted view is that the activation of A1/C1 neurons (and consequently AVP neurons) during hypoglycemia depends on peripheral glucose sensing at multiple sites, including the hepatic portal system (66, 67). We now show that activity of AVP neurons is increased when plasma glucose fall to ∼4.9 mM. The exact location of the glucose sensing is still controversial. It is unknown whether this threshold is sufficient to directly activate A1/C1 neurons. However, vagal afferents in the hepatic portal vein – which convey vagal sensory information to the A1/C1 neurons via projections to the nucleus of the solitary tract (66) - are responsive to alterations in glucose at this threshold (68).

In both mouse and human plasma, there was a relatively high basal level of copeptin. The high basal levels of copeptin likely relates to the longer half-life of this larger peptide (28, 29). Plasma AVP undergoes much more dramatic variations (28, 29), as our human data demonstrate but such measurements are currently not feasible in mice (26). We found that plasma AVP levels during hypoglycemia varies between 1 and 30 pM (**Figure 6c,d**), in good agreement with previous studies measuring AVP with a radioimmunoassay in human and rat (69, 70). This range of concentrations are also in line with the glucagonotrophic effect of AVP *in vitro* (**Figure 2e,g**) and the ability of AVP to increase alpha-cell membrane potential, [Ca^2+^]_i_, and [DAG]_i_ (**Figure 3**).

Cranial diabetes insipidus is a condition caused by reduced AVP production in the pituitary. Our findings provide an explanation to observations made 50 years ago that patients with cranial diabetes insipidus have a dramatically increased risk of hypoglycemia in response to the sulfonylurea chlorpropamide (71–73), with hypoglycemia occurring with prevalence of 50% and therefore considered the major barrier of this treatment (71–73). The present data raise the interesting possibility that this reflects the loss of AVP-induced glucagon secretion in these patients.

In T1D, deficiency of the secretory response of glucagon to hypoglycemia is an early acquired (< 2 years of onset) abnormality of counter-regulation and leads to severe hypoglycemia (74, 75), which may in part be explained by the major structural and functional changes that occur to islets in T1D. However, we also found that insulin-induced copeptin secretion was reduced, with some subjects exhibiting no elevation in copeptin. The reason for this change is unknown, but we speculate that recurrent hypoglycemia in patients with T1D may result in changes in glucose and/or insulin sensitivity in the A1/C1 region. Hypoglycemia awareness – a symptomatic component of which is driven by the autonomic nervous system – is associated with decreased insulin-induced copeptin in T1D patients (76). Our data suggest that monitoring of copeptin may prove an important tool for stratification of hypoglycemia risk in patients with type 1 diabetes.

## Acknowledgements

We would like to thank Prof. W. Scott Young and Emily Shepard from NIMH for kindly providing us with *Avpr1b* knockout mice and Professor Guy A. Rutter for hosting eye imaging experiments in his laboratory at Imperial College, UK.

We thank the Alberta Diabetes Institute IsletCore (University of Alberta, AB, Canada) for providing human islets, the isolation of which was supported in part by the Alberta Diabetes Foundation, the Human Organ Procurement and Exchange Program (Edmonton), and the Trillium Gift of Life Network (Toronto). We also thank Prof. Paul R.V. Johnson and colleagues at the Diabetes Research and Wellness Foundation Islet Isolation Facility. We would also like to thank the generosity of the organ donors and their families.

## Data availability statement

The authors declare that all data supporting the findings of this study are available within the article and its Supplementary Information or from the corresponding author on reasonable request.

## Author contributions

All authors collected and analysed the data. LJBB conceived the project and planned the experiments with AK. LJBB wrote the initial draft of the manuscript. LJBB and PR wrote the final version of the manuscript. All authors edited and approved the subsequent drafts of the manuscript. VS and KS designed and performed the *in vivo* alpha cell imaging experiments. The clinical experiments in human participants were conducted by MC and TM in the laboratory of FKK.

## Supplementary Figure Legends

**Supplementary Figure 1:**
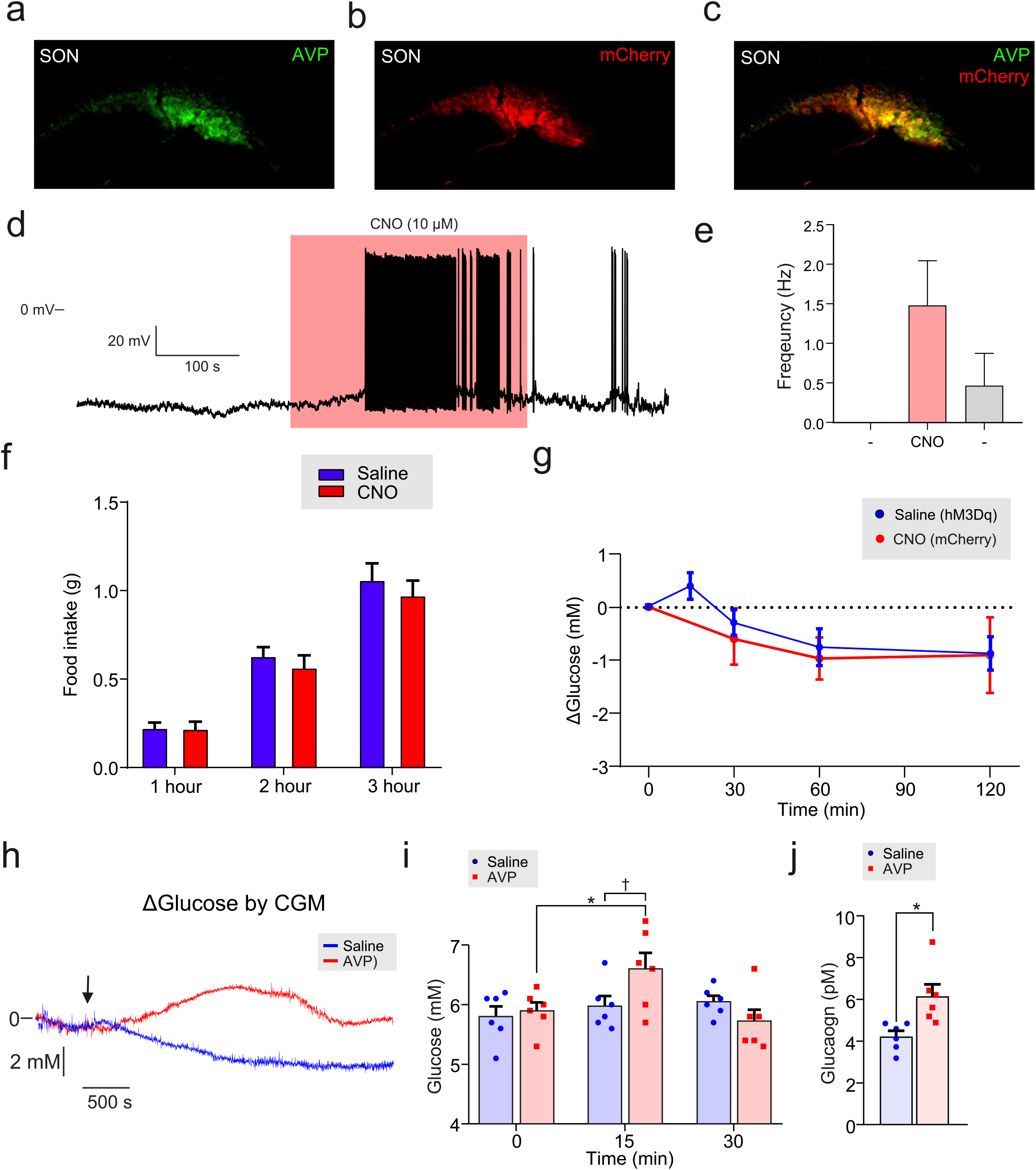
Effects of CNO in animals expressing mCherry in SON AVP neurons. (a) AAV-DIO-hM3Dq-mCherry was injected into the supraoptic nucleus (SON) of *Avp*^ires-Cre+^ mice. Immunostaining for AVP. See **Figure 1**. (b) Staining for mCherry; same brain sample as (a). (c) Merge of (a) and (b). (d) Patch-clamp recording from mCherry^+^ neuron in the SON in response to CNO. Firing frequency was increased in response to CNO. (e) Grouped data for firing frequency response to CNO in 3 mCherry^+^ neurons from SON of n=2 mice, as described in (a). One-way ANOVA; before treatment (-) vs. CNO, p=0.11. CNO concentration applied was 5-10 µM. (f) Food intake in response to CNO in *Avp*^ires-Cre+^ mice expressing hM3Dq in the SON. Two-way RM (both factors) ANOVA (Time, p<0.001; Treatment, p=0.52). n=11 mice. (g) AAV-DIO-mCherry was injected into the supraoptic nucleus (SON) of *Avp*^ires-Cre+^ mice, yielding mCherry expression in AVP neurons. CNO (3 mg/kg; i.p.) was injected and blood glucose measured (control experiments for **Figure 1b**). One-way RM ANOVA (Time, p=0.26). n=7 mice. For comparison, the saline injections from **Figure 1b** are also shown, wherein saline was injected i.p. into mice expressing hM3Dq in AVP neurons in the SON. (h) Response of blood glucose to exogenous AVP (5 µg/kg) during continuous glucose monitoring. Representative of 6 trials from n=2 mice. (i) AVP (10 µg/kg, i.p.) was injected into wild-type mice and blood glucose was measured with glucose test strips. Two-way RM ANOVA with Tukey’s (within treatments) and Sidak’s (between treatments) multiple comparison. AVP injection, 0 min vs 15 min, p<0.05=*. Vehicle, 0 mins vs 15 mins, p>0.2. Between treatments at t=15 mins; p=0.038, †. (j) Same cohort as in (e), but plasma glucagon measurements. Two-way RM ANOVA; p<0.05=*.

**Supplementary Figure 2:**
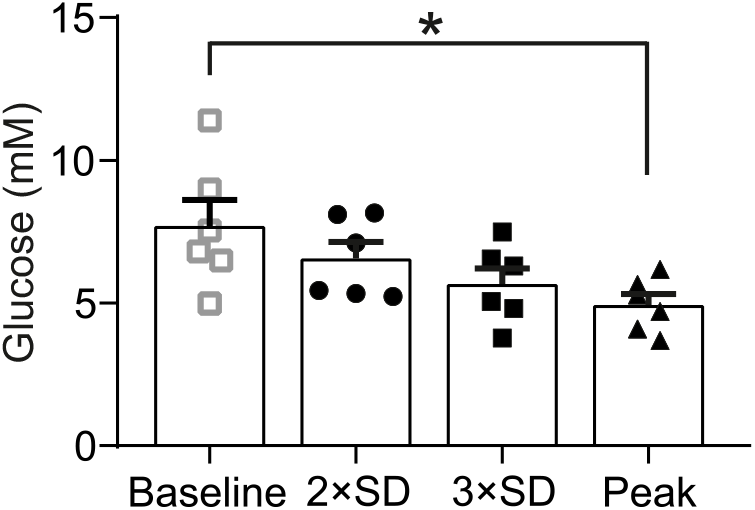
Simultaneous continuous glucose monitoring (CGM) and *in vivo* fiber photometry of AVP neurons. Grouped analysis of the glucose value at which the GCaMP6 signal crosses >2 SD from baseline, > 3 SD from baseline and first exhibits a peak. 6 trials, 2 mice. One-way repeated measures ANOVA (Sidak), p<0.05=*.

**Supplementary Figure 3:**
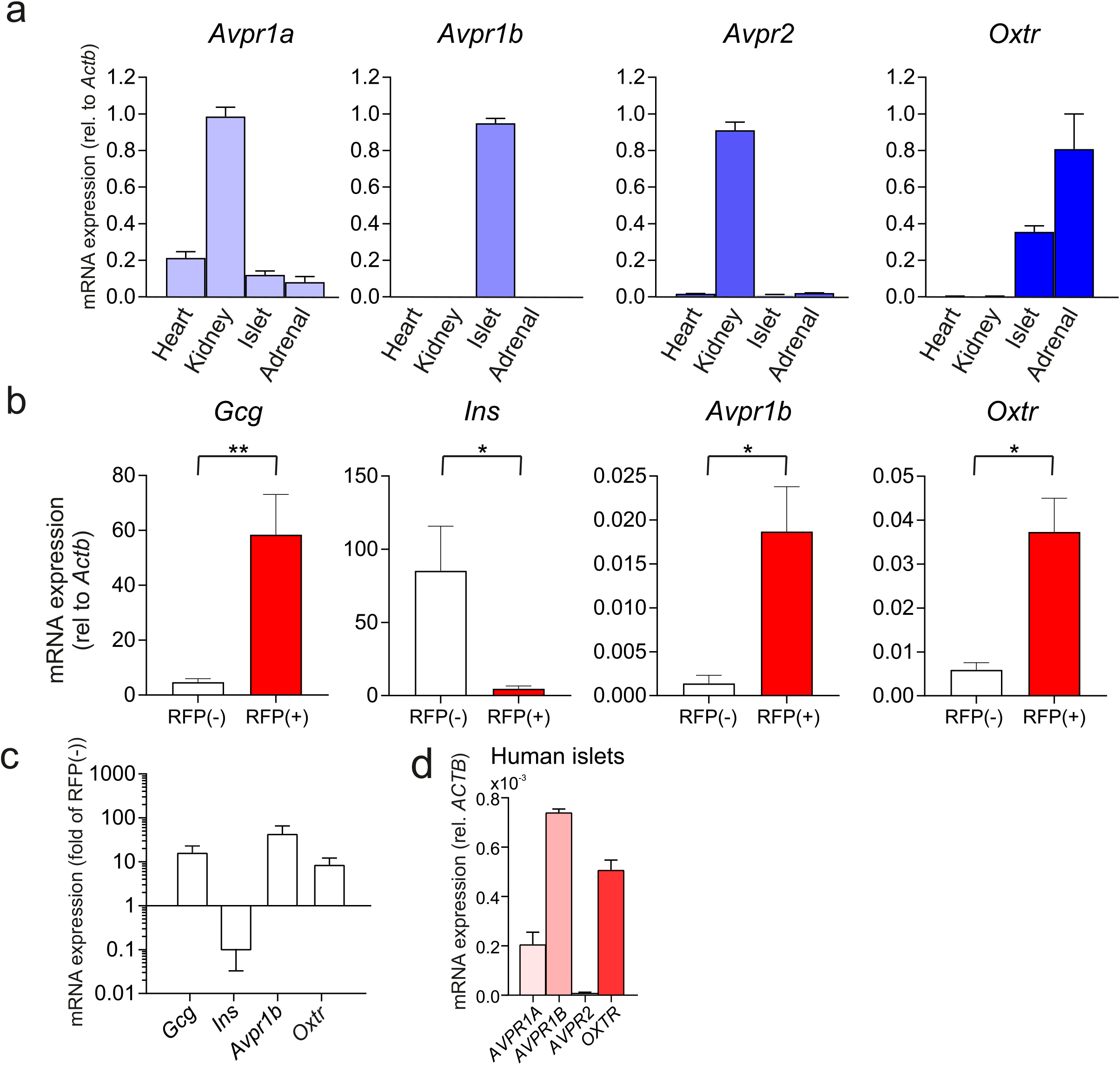
Expression of the vasopressin 1b receptor in mouse and human. (a) mRNA expression of *Avpr* family in mouse heart, kidney, islets and adrenal glands. Samples from n=3 wild-type mice, each run in triplicate. Calculated with the Pfaffl method, using *Actb* as the reference gene. (b) mRNA expression in sorted islet cells. Fractions were sorted from mice with a fluorescent reporter (RFP) in alpha-cells (*Gcg*^Cre+^-RFP mice) into an alpha-cell fraction (RFP(+)) and non-alpha-cell fraction (RFP(-)). Data from 4 sorts from n=4 *Gcg*^Cre+^-RFP mice. Ratio paired t-test; p<0.05=*, p<0.01=**. (c) Same data as in b). mRNA expression in alpha-cells (RFP(+) fraction) represented as fold of RFP(-) expression. Scale = log_10_. All data represented as mean ± SEM. (d) mRNA expression of *AVPR1A*, *AVPR1B*, AVPR2 and *OXTR* in human islets. Human islet samples n=7-8 donors. Calculated with the Pfaffl method, using *Actb* as the reference gene.

**Supplementary Figure 4:**
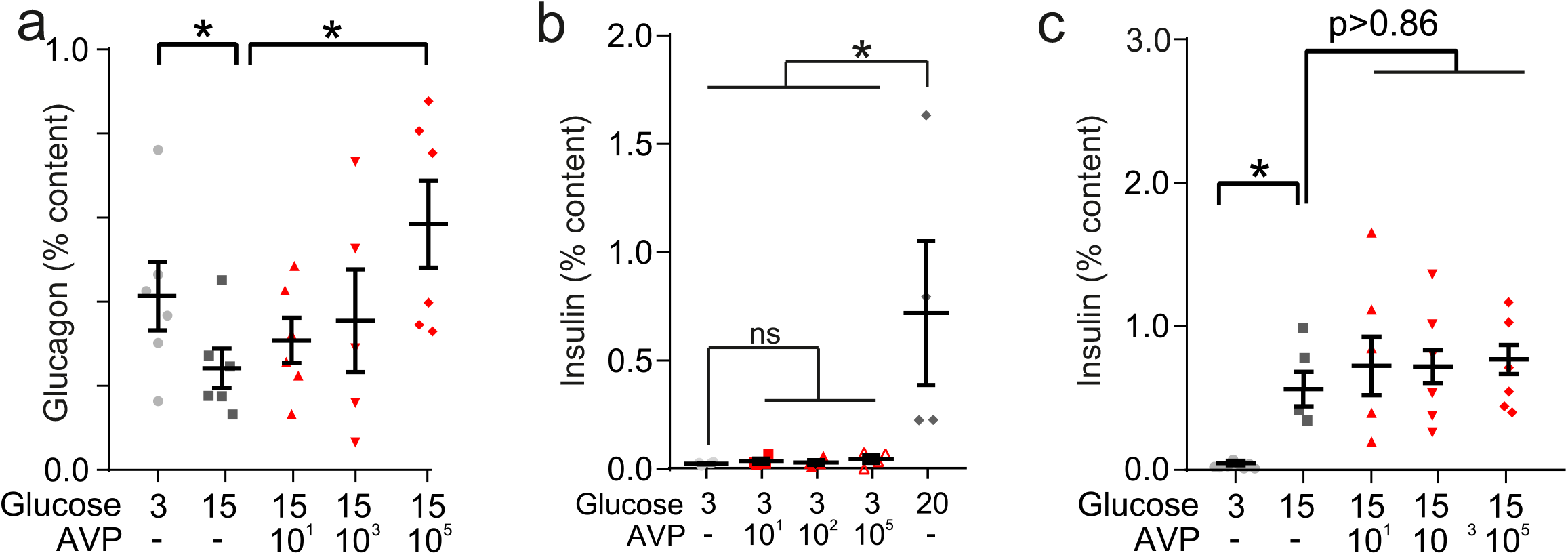
AVP and beta-cell function. (a) Glucagon secretion (% content) from isolated mouse islets in response to AVP. One-way ANOVA (p<0.05=*). n=5-6 wild-type mice per condition. AVP was applied in high (15 mM glucose). (b) Insulin secretion from isolated mouse islets in response to physiological (10-100 pM) and supra-physiological (100 nM) concentrations of AVP. One-way ANOVA, p<0.01=**. n=5 wild-type mice per condition. (c) Insulin secretion (% content) from isolated mouse islets in response to AVP. One-way ANOVA, p<0.01=**. n=4-6 wild-type mice per condition. AVP was applied in high (15 mM glucose).

**Supplementary Figure 5:**
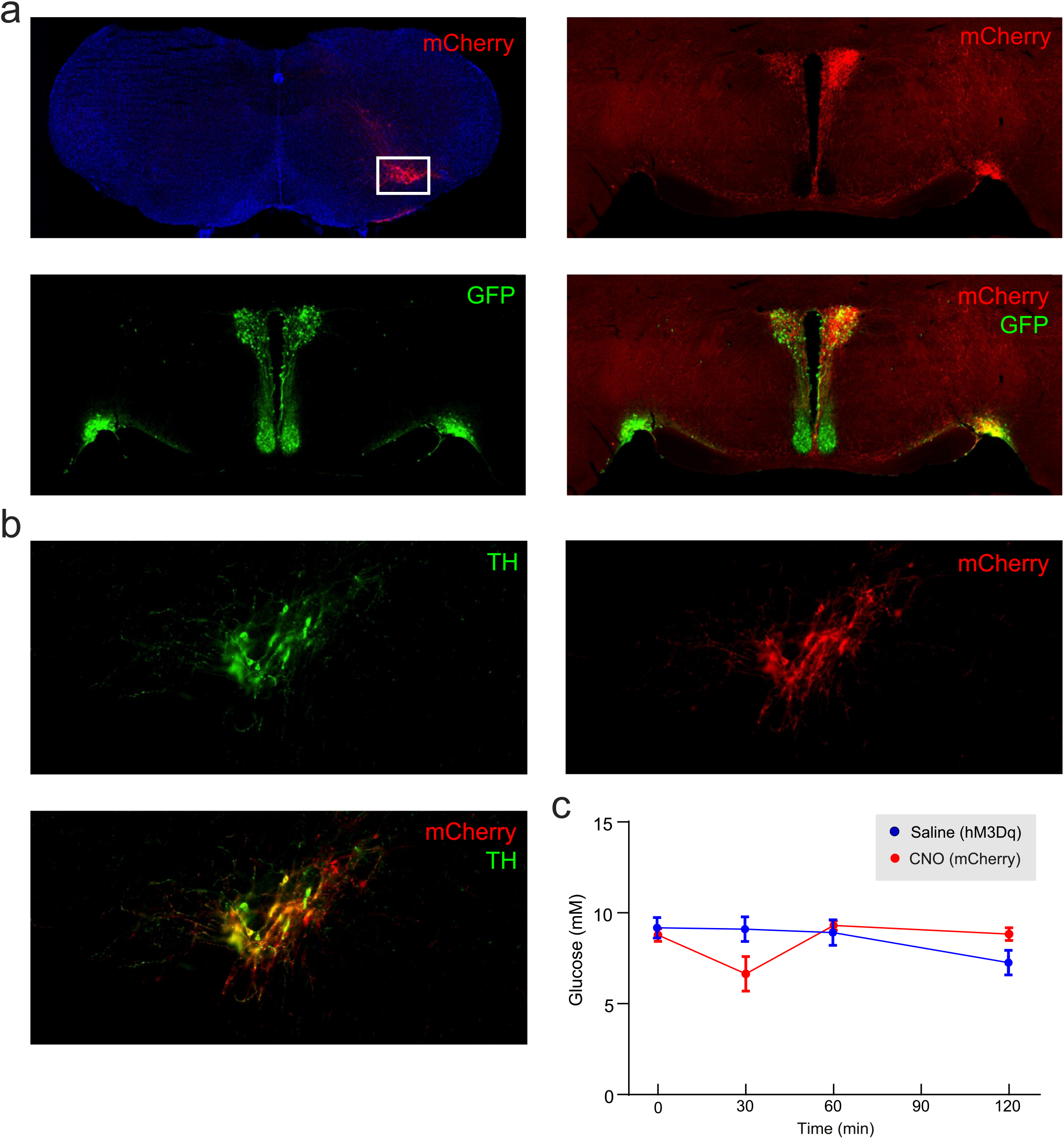
Viral tracing of A1/C1 terminals. (a) Injection of a Cre-dependent viral vector containing the light-gated ion channel Channelrhodopsin-2 (AAV-DIO-ChR2-mCherry) into A1/C1 neurons of *Th*^Cre+^ mice. *Top left:* mCherry expression in A1/C1 neurons (white box). *Top right:* mCherry expression in the PVH and SON. *Bottom left:* AVP-immunoreactive neurons (green) expression in the PVH and SON. *Bottom right:* A1/C1 terminals (mCherry) co-localizing with AVP neurons (green) in the PVH and SON. (b) Expression of hM3Dq in A1/C1 neurons, following AAV-DIO-hM3Dq-mCherry injection into the A1/C1 region of *Th*^Cre+^ mice (i). TH = tyrosine hydroxylase. (c) CNO administration (i.p.) into *Th*^Cre+^ mice expressing mCherry (and not hM3Dq) in A1/C1 neurons (control experiments for **Figure 5f-h**). The saline injections in mice expressing hM3Dq in A1/C1 neurons (from **Figure 5f-h**) are also shown for comparison.

**Supplementary Figure 6:**
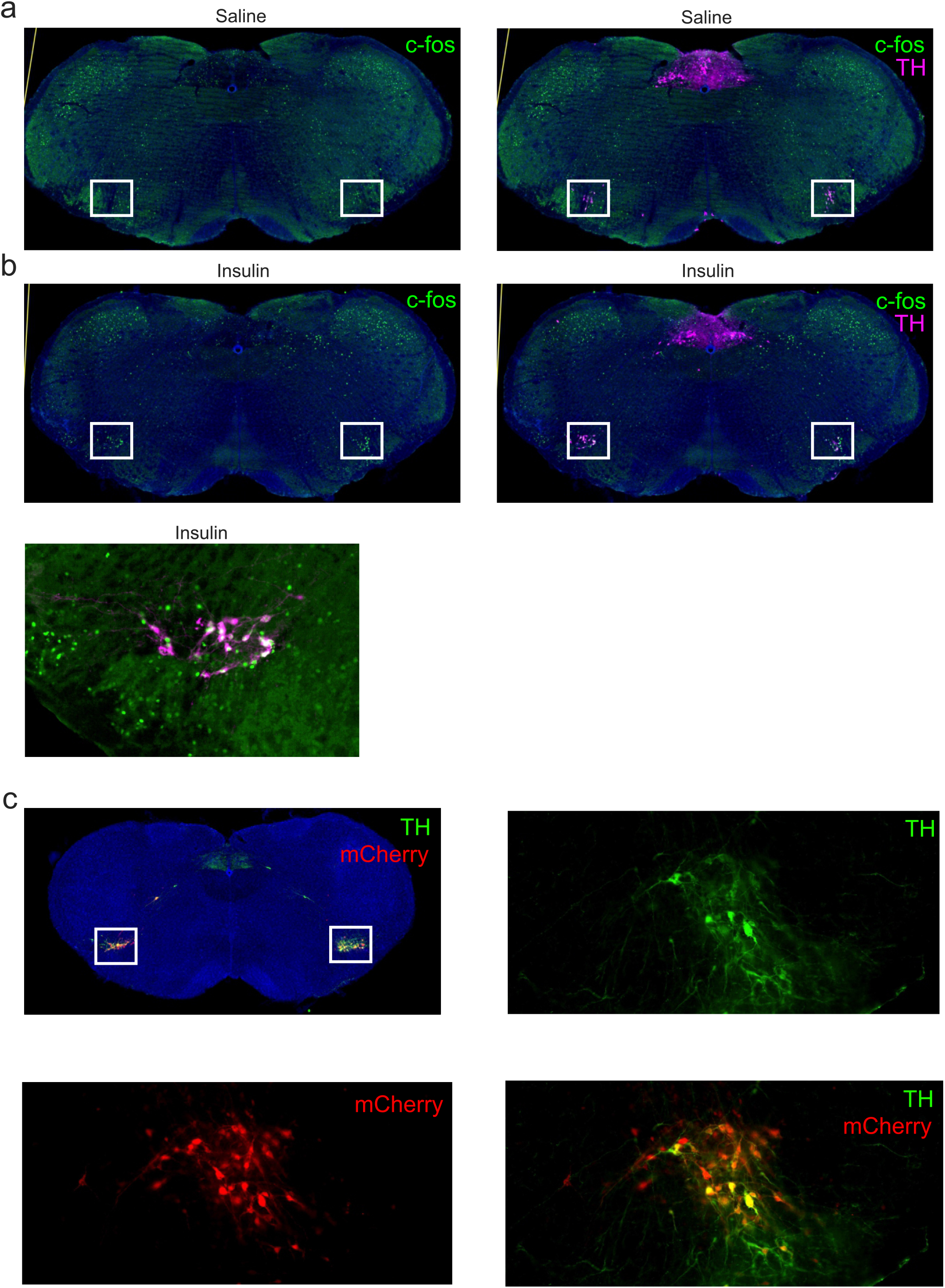
c-fos expression in A1/C1 neurons during an ITT. (a) *Left: c-fos* expression 30 minutes following i.p. saline injection. *Right:* Same animal, but merge of *Th* and *c-fos*. Note the *Th^+^* immunoreactive neurons in the A1/C1 region (white box). (b) *Left: c-fos* expression in *Th*^Cre+^ mice following an 30 minutes following i.p. insulin injection. *Right:* Same animal, but merge of *Th* and *c-fos*. Note the *Th^+^* immunoreactive neurons in the A1/C1 region (white box) co-expressing *c-fos*. *Lower panel:* Magnified view of *c-fos* and *Th* in A1/C1 neurons 30 minutes following insulin injection. Images representative of experiments in n=1+1 mice. See **Figure 5**. (c) *Top left:* Expression of mCherry and TH (green) in A1/C1 neurons (white box) of *Dbh*^flp+^ mice, following viral injection of AAV-fDIO-hM4Di-mCherry into the VLM, targeting A1/C1 neurons. *Top right:* Magnified view of the A1/C1 region, TH expression. *Bottom left:* Magnified view, mCherry expression. *Bottom right:* merge. TH = tyrosine hydroxylase.

**Supplementary Table 1:** Participant characteristics for Figure 7.

|  | Christensen *et al.* (2011) | Christensen *et al.* (2015) | p |
| --- | --- | --- | --- |
| Cohort | Control (n=10) | T1D (n=10) |  |
| Age (years) | 23±1 | 26±1 | 0.06 |
| BMI (kg/m<sup>2</sup>) | 23±0.5 | 24±0.5 | 0.1744 |
| HbA1c (%) | 5.5±0.1 | 7.3±0.2 | <0.0001 |

## Methods

### Ethics

All animal experiments were conducted in strict accordance to regulations enforced by the research institution. Experiments conducted in the UK were done so in accordance with the UK Animals Scientific Procedures Act (1986) and University of Oxford and Imperial College London ethical guidelines, and were approved by the local Ethical Committee. All animal care and experimental procedures conducted in the U.S.A. were approved by the Beth Israel Deaconess Medical Center Institutional Animal Care and Use Committee. Animal experiments conducted in Goteborg University were approved by a local Ethics Committee.

Human pancreatic islets were isolated, with ethical approval and clinical consent, at the Diabetes Research and Wellness Foundation Human Islet Isolation Facility (OCDEM, Oxford, UK) or Alberta Diabetes Institute IsletCore (University of Alberta, AB, Canada). Islets from a total of 19 human donors were used in this study. Donor details were as follows; age = 42 ± 4 years; BMI = 27.1 ± 3; Sex = 10/9 (M/F).

### Animals

All animals were kept in a specific pathogen-free (SPF) facility under a 12:12 hour light:dark cycle at 22 °C, with unrestricted access to standard rodent chow and water. C57BL/6J mice used in this study are referred to as wild-type mice. To generate alpha-cell specific expression of the genetically-encoded Ca^2+^ sensor GCaMP3, mice carrying Cre recombinase under the control of the proglucagon promoter (*Gcg*^Cre+^ mice) were crossed with mice with a floxed green calmodulin (GCaMP3) Ca^2+^ indicator in the ROSA26 locus (The Jackson Laboratory). These mice are referred to as *Gcg*-GCaMP3 mice. To generate mice expressing RFP in alpha-cells, *Gcg*^Cre+^ were crossed with mice containing a floxed tandem-dimer red fluorescent protein (tdFRP) in the ROSA26 locus (*Gcg*-RFP mice). Both of these mouse models were kept on a C57BL/6J background. Other transgenic mouse strains used – namely, *Avp*^ires-Cre+^ (77), *Th*^Cre+^ (The Jackson Laboratory), *Dbh*^flp+^ (MMRCC) and *Avp*^GFP^ (MMRCC) - were heterozygous for the transgene and maintained on a mixed background. *Avpr1b*^−/−^ and littermate controls (*Avpr1b^+/+^*) were bred and maintained as previously described (78).

### Isolation of mouse islets

Mice of both sex and 11-16 weeks of age were killed by cervical dislocation (UK Schedule 1 procedure). Pancreatic islets were isolated by liberase digestion followed by manual picking. Islets were used acutely and were, pending the experiments, maintained in tissue culture for <36 hour in RPMI 1640 (11879-020, Gibco, Thermo Fisher Scientific) containing 1% pennicillin/streptomycin (1214-122, Gibco, Thermo Fisher Scientific), 10%FBS (F7524-500G, Sigma-Aldrich) and 11 mM glucose, prior to experiments.

### Patch-clamp electrophysiology in islets

Mouse islets were used for patch-clamp electrophysiological recordings. These recordings (in intact islets) were performed at 33-34 °C using an EPC-10 patch-clamp amplifier and PatchMaster software (HEKA Electronics, Lambrecht/Pfalz, Germany). Unless otherwise stated, recordings were made in 3 mM glucose, to mimic hypoglycemic conditions in mice. Currents were filtered at 2.9 kHz and digitized at > 10 kHz. A new islet was used for each recording. Membrane potential (*V*_*M*_) recordings were conducted using the perforated patch-clamp technique, as previously described (9). The pipette solution contained (in mM) 76 K_2_SO_4_, 10 NaCl, 10 KCl, 1 MgCl_2_·6H_2_0 and 5 Hepes (pH 7.35 with KOH). For these experiments, the bath solution contained (mM) 140 NaCl, 3.6 KCl, 10 Hepes, 0.5 MgCl_2_·6H20, 0.5 Na_2_H_2_PO_4,_ 5 NaHCO_3_ and 1.5 CaCl_2_ (pH 7.4 with NaOH). Amphotericin B (final concentration of 25 mg/mL, Sigma-Aldrich) was added to the pipette solution to give electrical access to the cells (series resistance of <100 MΩ). Alpha-cells in *Gcg*-GCaMP3 islets were confirmed by the presence of GCaMP3.

The frequency of action potential firing was calculated in MATLAB v. 6.1 (2000; The MathWorks, Natick, MA). In brief, a peak-find algorithm was used to detect action potentials. This was then used to calculate firing frequency in different experimental conditions (AVP concentrations). Power-spectrum analysis of *V*_*M*_ was conducted in Spike2 (CED, Cambridge, UK). *V*_*M*_ was moving-average filtered (interval of 200 ms) and the mean *V*_*M*_ subtracted. A power-spectrum was then produced (Hanning window with 0.15 Hz resolution) during 3 mM glucose alone, and with 10 pM AVP.

### GCaMP3 imaging in mouse islets

Time-lapse imaging of the intracellular Ca^2+^ concentration ([Ca^2+^]_i_) in *Gcg*-GCaMP3 mouse islets was performed on an inverted Zeiss AxioVert 200 microscope, equipped with the Zeiss LSM 510-META laser confocal scanning system, using a 403/1.3 NA objective. Mouse islets were transferred into a custom-built recording chamber. Islets were then continuously perfused with bath solution at a rate of 200 µL/min. The bath solution contained (in mM): 140 NaCl, 5 KCl, 1.2 MgCl_2_, 2.6 CaCl_2_, 1 NaH_2_PO_4_, 10 Hepes, 17 mannitol and 3 glucose. GCaMP3 was excited at 430 nm and recorded at 300-530 nm. The pinhole diameter was kept constant, and frames of 256 x 256 pixels were taken every 800 ms. Unless otherwise stated, recordings were made in 3 mM glucose, to mimic hypoglycemic conditions in mice. Raw GCaMP3 data was processed as follows; regions of interest (ROIs) were manually drawn around each GCaMP3^+^ cell in ImageJ and the time-series of the GCaMP3 signal for each cell was exported. These data were first imported into Spike2 7.04 (CED, Cambridge, UK), wherein the data was median filtered to remove baseline drift. The size of the filter was optimised for each individual cell to remove drift/artefacts but preserve Ca^2+^ transients. Ca^2+^ transients were then automatically detected using the built in peak-find algorithm; the amplitude of peaks to be detected was dependent on the SNR but was typically > 20% of the maximal signal intensity. Following this, frequency of Ca^2+^ transients could be determined. For plotting Ca^2+^ data, the data was imported into MATLAB.

### DAG measurements in mouse islets

The effects of AVP on the intracellular diacylglycerol concentration (DAG) in pancreatic islet cells was studied using a recombinant circularly permutated probe, Upward DAG (Montana Molecular). Islets isolated from wild-type mice were gentle dispersed (using Trypsin ES) into clusters and platted on rectangular coverslips. Cell clusters were then transfected with Upward DAG, delivered via a BacMam infection (according to the manufacturer’s guidelines). Coverslips were then were placed in a custom built chamber. Imaging experiments were performed 36-48 hours after infection using a Zeiss AxioZoom.V16 zoom microscope equipped with a ×2.3/0.57 objective (Carl Zeiss). The fluorescence was excited at 480 nm, and the emitted light was collected at 515 nm. The cells were kept at 33-35 °C and perfused continuously throughout the experiment with KRB solution supplemented with 3 mM glucose. The images were acquired using Zen Blue software (Carl Zeiss). The mean intensity for each cell was determined by manually drawing ROIs in ImageJ. Data analysis and representation was performed with MATLAB. All data was processed using a moving average filter function (*smooth*) with a span of 50 mins, minimum subtracted and then normalised to maximum signal intensity in the time-series. AUC was calculated using the *trapz* function and then divided by the length of the condition.

### Ca^2+^ imaging in human islets

Time-lapse imaging of [Ca^2+^]_i_ in human islets was performed on the inverted Zeiss SteREO Discovery V20 Microscope, using a PlanApo S 3.5x mono objective. Human islets were loaded with 5 µg/µL of the Ca^2+^-sensitive dye Fluo-4 (1923626, Invitrogen, Thermo Fisher Scientific) for 60 min before being transferred to a recording chamber. Islets were then continuously perfused with DMEM (11885-084, Gico, Thermo Fisher Scientific) with 10% FBS, 100 units/mL penicillin and 100 mg/mL streptomycin at a rate of 200 µL/min. Fluo-4 was excited at 488 nm and fluorescence emission collected at 530 nm. The pinhole diameter was kept constant, and frames of 1388×1040 pixels were taken every 3 sec. The mean intensity for each islet was determined by manually drawing an ROI around the islet in ImageJ. Data analysis and representation was performed with MATLAB. All data was processed using a moving average filter function (*smooth*) with a span of 20 mins, minimum subtracted and then normalised to maximum signal intensity in the time-series. AUC was calculated using the *trapz* function and then divided by the length of the condition.

### Pancreatic islet isolation, transplantation and *in vivo* imaging of islets implanted into the anterior chamber of the eye (ACE)

Pancreatic islets from *Gcg*-GCaMP3 mice were isolated and cultured as described above. For transplantation, 10-20 islets were aspirated with a 27-gauge blunt eye cannula (BeaverVisitec, UK) connected to a 100 µl Hamilton syringe (Hamilton) via 0.4-mm polyethylene tubing (Portex Limited). Prior to surgery, mice (C57BL6/J) were anesthetised with 2-4% isoflurane (Zoetis) and placed in a stereotactic frame. The cornea was incised near the junction with the sclera, then the blunt cannula (pre-loaded with islets) was inserted into the ACE and islets were expelled (average injection volume 20 μl for 10 islets). Carprofen (Bayer, UK) and eye ointment were administered post-surgery. A minimum of four weeks was allowed for full implantation before imaging. Imaging sessions were performed with the mouse held in a stereotactic frame and the eye gently retracted, with the animal maintained under 2-4% isoflurane anesthesia. All imaging experiments were conducted using a spinning disk confocal microscope (Nikon Eclipse Ti, Crest spinning disk, 20x water dipping 1.0 NA objective). The signal from GCaMP3 (ex. 488 nm, em. 525±25 nm) was monitored at 3 Hz for up to 20 min. After a baseline recording, mice received a bolus of AVP (10 µg/kg) i.v. (tail vein). Data were imported into ImageJ for initial movement correction (conducted with the StackReg and TurboReg plugins) and ROI selection. Analysis was then conducted in MATLAB.

### Hormone secretion measurements from mouse and human islets

Islets, from human donors or isolated from wild-type mice, were incubated for 1 h in RPMI or DMEM supplemented with 7.5 mM glucose in a cell culture incubator. Size-matched batches of 15-20 islets were pre-incubated in 0.2 ml KRB with 2 mg/ml BSA (S6003, Sigma-Aldrich) and 3 mM glucose for 1 hour in a water-bath at 37 °C. Following this islets were statically subjected to 0.2 ml KRB with 2 mg/ml BSA with the condition (e.g. 10 pM AVP) for 1 hour. After each incubation, the supernatant was removed and kept, and 0.1 ml of acid:etoh (1:15) was added to the islets. Both of these were then stored at −80 °C. Each condition was repeated in at least triplicates.

Glucagon and insulin measurements in supernatants and content measurements were performed using a dual mouse insulin/glucagon assay system (Meso Scale Discovery, MD, U.S.A.) according to the protocol provided.

### Hormone secretion measurements in the perfused mouse pancreas

Dynamic measurements of glucagon were performed using the *in situ* perfused mouse pancreas. Briefly, the aorta was cannulated by ligating above the coeliac artery and below the superior mesenteric artery, and the pancreas was perfused with KRB at a rate of ∼0.45ml/min using an Ismatec Reglo Digital MS2/12 peristaltic pump. The KRB solution was maintained at 37 °C with a Warner Instruments temperature control unit TC-32 4B in conjunction with a tube heater (Warner Instruments P/N 64-0102) and a Harvard Apparatus heated rodent operating table. The effluent was collected by cannulating the portal vein and using a Teledyne ISCO Foxy R1 fraction collector. The pancreas was first perfused for 10 min with 3 mM glucose before commencing the experiment to establish the basal rate of secretion. Glucagon measurements in collected effluent were performed using RIA.

### Flow cytometry of islet cells (FACS), RNA extraction, cDNA synthesis and quantitative PCR

The expression of the AVPR gene family was analysed in tissues from 12-week old C57BL6/J mice (3 mice) and pancreatic islets from human donors (2 samples, each comprised of pooled islet cDNA from 7 and 8 donors, respectively). Total RNA was isolated using a combination of TRIzol and PureLink RNA Mini Kit (Ambion, Thermofisher Scientific) with incorporated DNase treatment.

Pancreatic islets from *Gcg*-RFP mice were isolated and then dissociated into single cells by trypsin digestion and mechanical dissociation. Single cells were passed through a MoFlo Legacy (Beckman Coulter). Cells were purified by combining several narrow gates. Forward and side scatter were used to isolate small cells and to exclude cell debris. Cells were then gated on pulse width to exclude doublets or triplets. RFP^+^ cells were excited with a 488 nm laser and the fluorescent signal was detected through a 580/30 bandpass filter (*i.e*. in the range 565-595 nm). RFP-negative cells were collected in parallel. The levels of gene expression in the RFP^+^ and in the RFP^−^ FAC-sorted fractions were determined using real-time quantitative PCR (qPCR). RNA from FACS-sorted islet cells was isolated using RNeasy Micro Kit (Qiagen). cDNA was synthesized using the High Capacity RNA-to-cDNA kit (Applied Biosystems, Thermofisher Scientific). Real time qPCR was performed using SYBR Green detection and gene specific QuantiTect Primer Assays (Qiagen) on a 7900HT Applied Biosystems analyser. All reactions were run in triplicates. Relative expression was calculated using ΔCt method *Actb* as a reference gene.

### Stereotaxic surgery and viral injections

For viral injections into the SON, mice were anesthetized with ketamine/xylazine (100 and 10 mg/kg, respectively, i.p.) and then placed in a stereotaxic apparatus (David Kopf model 940). A pulled glass micropipette (20-40 μm diameter tip) was used for stereotaxic injections of adeno-associated virus (AAV). Virus was injected into the SON (200 nl/side; AP: −0.65 mm; ML: ±1.25 mm; DV: −5.4 mm from bregma) by an air pressure system using picoliter air puffs through a solenoid valve (Clippard EV 24VDC) pulsed by a Grass S48 stimulator to control injection speed (40 nL/min). The pipette was removed 3 min post-injection followed by wound closure using tissue adhesive (3M Vetbond). For viral injections into the VLM, mice were placed into a stereotaxic apparatus with the head angled down at approximately 45°. An incision was made at the level of the cisterna magna, then skin and muscle were retracted to expose the dura mater covering the 4th ventricle. A 28-gauge needle was used to make an incision in the dura and allow access to the VLM. Virus was then injected into the VLM (50nl*2/side; AP: −0.3 and −0.6mm; ML: ±1.3mm; DV: −1.7mm from obex) as described above. The pipette was removed 3 min post-injection followed by wound closure using absorbable suture for muscle and silk suture for skin. For fiber photometry, an optic fiber (200 µm diameter, NA=0.39, metal ferrule, Thorlabs) was implanted in the SON and secured to the skull with dental cement. Subcutaneous injection of sustained release Meloxicam (4 mg/kg) was provided as postoperative care. The mouse was kept in a warm environment and closely monitored until resuming normal activity. Chemogenetic experiments utilized AAV8-hSyn-DIO-hM3Dq-mCherry (Addgene:44361) and AAV8-nEF-fDIO-hM4Di-mCherry (custom-made vector) produced from Boston Children’s Hospital Viral Core and AAV5-EF1α-DIO-mCherry purchased from the UNC Vector Core. Fiber photometry experiments were conducted using AAV1-hSyn-FLEX-GCaMP6s purchased from the University of Pennsylvania (School of Medicine Vector Core). Projection mapping and ChR2-assisted circuit mapping were done using AAV9-EF1α-DIO-ChR2(H134R)-mCherry purchased from the University of Pennsylvania (School of Medicine Vector Core).

### Fiber photometry experiments and analysis of photometry data

*In vivo* fiber photometry was conducted as previously described (Mandelblat-Cerf, et al. (79)). A fiber optic cable (1-m long, metal ferrule, 400 µm diameter; Doric Lenses) was attached to the implanted optic cannula with zirconia sleeves (Doric Lenses). Laser light (473 nm) was focused on the opposite end of the fiber optic cable to titrate the light intensity entering the brain to 0.1-0.2 mW. Emitted light was passed through a dichroic mirror (Di02-R488-25×36, Semrock) and GFP emission filter (FF03-525/50-25, Semrock), before being focused onto a sensitive photodetector (Newport part #2151). The GCaMP6 signal was passed through a low-pass filter (50 Hz), and digitized at 1 KHz using a National Instruments data acquisition card and MATLAB software.

All experiments were conducted in the home-cage in freely moving mice. Animals prepared for *in vivo* fiber photometry experiments (outlined above), were subjected to an ITT or 2DG injection after overnight fasting. Prior to insulin or 2DG injection, a period of GCaMP6s activity was recorded (3 min) to establish baseline activity. Insulin (i.p. 2 U/kg), 2DG (i.p. 500mg/kg), or saline vehicle was then administered, and GCaMP6 activity recorded for a further 40 min. In some experiments, mice were pre-treated i.p. with CNO (1mg/kg) or saline, 30 minutes prior to insulin or 2DG. The recorded data was exported and then imported into MATLAB for analysis. Fluorescent traces were down-sampled to 1 Hz and the signal was normalised to the baseline (F_0_ mean activity during baseline activity), with 100% signal being defined as the maximum signal in the entire trace (excluding the injection artefact). Following the ITT, the signal was binned (1 min) and a mean for each bin calculated. These binned signals were compared to baseline signal using a one-way RM ANOVA.

### Surgery for continuous glucose monitoring

Animals that have undergone fiber photometry surgeries (3 weeks prior) were anesthetized and maintained with isoflurane. Once mice were fully anesthetized, the ventral abdomen and underside of the neck were shaved and disinfected. Animals were placed on their backs on a heated surgical surface. For transmitter implantation, a ventral midline abdomen incision was made and the abdominal wall was incised. The transmitter was placed in the abdominal cavity with the lead exiting cranially and the sensor and connector board exteriorized. The incision was sutured incorporating the suture rib into the closure. For glucose probe implantation, a midline neck incision was performed and the left common carotid artery was isolated. The vessel was then perforated and the sensor of the glucose probe (HD-XG, Data Sciences International) was advanced into the artery towards the heart, within a final placement in the aortic arch. Once in place, the catheter was secured by tying the suture around the catheter and vessel, and overlying opening in tissue was closed. Mice were kept warm on a heating pad and monitored closely until fully recovered from anesthesia.

### Simultaneous AVP fiber photometry and continuous glucose monitoring

All experiments were conducted in the home-cage in freely moving mice. Animals prepared for in vivo fiber photometry and continuous glucose monitoring (outlined above), were subjected to an ITT after overnight fasting. After establishing > 3 min of baseline activity, insulin (i.p. 1 or 1.5 U/kg) or saline vehicle was administered. GCaMP6s activity and blood glucose were recorded throughout 2 h of experiment. Each recording was separated by at least 48h. GCaMP6s recording was performed as described above. Blood glucose was acquired using Dataquest A.R.T. 4.36 system and analysed using MATLAB. Calibration of HD-XG device was performed as per manufacturer’s manual.

### *In vivo* measurements of plasma glucose, glucagon and copeptin

Samples for blood glucose and plasma glucagon measurements were taken from mice in response to different metabolic challenges (described in detail below). Both sexes were used for these experiments. Blood glucose was measured with an Accu-Chek Aviva (Roche Diagnostic, UK) and OneTouch Ultra (LifeScan, UK). Plasma copeptin in mouse was measured using an ELISA (MyBioSource, USA and Neo Scientific, USA). We note that the Kryptor BRAHMs system used for human samples could not be used for mouse samples (minimum sample volume of 250 µL plasma).

#### Insulin tolerance test

Mice were restrained and a tail vein sample of blood was used to measure fed plasma glucose. A further sample was extracted into EDTA coated tubes for glucagon measurements. Aprotinin (1:5, 4 TIU/ml; Sigma-Aldrich, UK) was added to all blood samples. These blood samples were kept on ice until the end of the experiment. Mice were first administered with any necessary pre-treatment and then individually caged. Pre-treatments included SSR149415 (30 mg/kg in PBS with 5% DMSO and 5% Cremophor EL), LY2409021 (5 mg/kg in PBS with 5% DMSO), CNO (1-3 mg/kg in PBS with 5% DMSO) or the appropriate vehicle. After a 30 minute period, mice were restrained again, and blood was taken via a tail vein or submandibular bleed. This was used for blood glucose measurements, and also for glucagon. Insulin (0.75, 1 or 1.5 U/kg) was then administered i.p., and the mice were re-caged. At regular time intervals after the insulin injection, mice were restrained and a blood sample extracted. Blood glucose was measured, and blood was taken for glucagon measurements. At the end of the experiment, blood samples were centrifuged at 2700 rpm for 10 min at 4 °C to obtain plasma. The plasma was then removed and stored at −80 °C. Plasma glucagon measurements were conducted using the 10-µl glucagon assay system (Mercodia, Upsala, Sweden), according to the manufacturer’s protocol.

#### Glucoprivic response to 2-Deoxy-D-glucose

Wild-type mice were used for 2-Deoxy-D-glucose (2DG) experiments. The mice were single housed one week prior to experimental manipulation. On the experimental day, food was removed 4 hours prior to the experiment. 2DG (500 mg/kg) or saline vehicle was then administered i.p., and blood samples taken at regular intervals for blood glucose and plasma glucagon measurements. In some cohorts, the V1bR antagonist SSR149415 (30 mg/kg in PBS with 5% DMSO and 5% Tween 80), glucagon receptor antagonist LY240901 (5 mg/kg in PBS with 5% DMSO and 5% Tween 80) or appropriate vehicle was administered i.p. 30 minutes prior to administration of 2DG. Plasma glucagon was measured as described above.

### Brain slice electrophysiology

To prepare brain slices for electrophysiological recordings, brains were removed from anesthetized mice (4–8 weeks old) and immediately placed in ice-cold cutting solution consisting of (in mM): 72 sucrose, 83 NaCl, 2.5 KCl, 1 NaH_2_PO_4_, 26 NaHCO_3_, 22 glucose, 5 MgCl_2_, 1 CaCl_2_, oxygenated with 95% O_2_ /5% CO_2_, measured osmolarity 310-320 mOsm/l. Cutting solution was prepared and used within 72 hours. 250 μm-thick coronal sections containing the PVH and SON were cut with a vibratome (7000smz2-Campden Instruments) and incubated in oxygenated cutting solution at 34 °C for 25 min. Slices were transferred to oxygenated aCSF (126 mM NaCl, 21.4 mM NaHCO_3_, 2.5 mM KCl, 1.2 mM NaH_2_PO_4_, 1.2 mM MgCl_2_, 2.4 mM CaCl_2_, 10 mM glucose) and stored in the same solution at room temperature (20-24 °C) for at least 60 min prior to recording. A single slice was placed in the recording chamber where it was continuously super-fused at a rate of 3–4 ml per min with oxygenated aCSF. Neurons were visualized with an upright microscope equipped with infrared-differential interference contrast and fluorescence optics. Borosilicate glass microelectrodes (5–7 MΩ) were filled with internal solution. All recordings were made using Multiclamp 700B amplifier, and data was filtered at 2 kHz and digitized at 10 kHz. All analysis was conducted off-line in MATLAB.

#### Channelrhodopsin-2 assisted circuit mapping (CRACM) of connections from A1/C1 neurons to the SON

A Cre-dependent viral vector containing the light-gated ion channel channelrhodopsin-2 (AAV-DIO-ChR2-mCherry) was injected into the VLM (targeting A1/C1 neurons) of *Avp*^GFP^ x *Th*^Cre+^ mice (see **Figure 5a**) as described above (see ‘Stereotaxic surgery and viral injections’). Brain slices were prepared (as above) from these mice. The SON was located by using the bifurcation of the anterior and middle cerebral arteries on the ventral surface of the brain as a landmark. Sections of 250 µm thickness of the SON were then cut with a Leica VT1000S or Campden Instrument 7000smz-2 vibratome, and incubated in oxygenated aCSF (126 mM NaCl, 21.4 mM NaHCO_3_, 2.5 mM KCl, 1.2 mM NaH_2_PO_4_, 1.2 mM MgCl_2_, 2.4 mM CaCl_2_, 10 mM glucose) at 34 °C for 25 min. Slices recovered for 1 hr at room temperature (20–24°C) prior to recording. Whole-cell voltage clamp recordings were obtained using a Cs-based internal solution containing (in mM): 135 CsMeSO_3_, 10 HEPES, 1 EGTA, 4 MgCl_2_, 4 Na_2_-ATP, 0.4 Na_2_-GTP, 10 Na_2_-phosphocreatine (pH 7.3; 295 mOsm). To photostimulate ChR2-positive A1/C1 fibers, an LED light source (473 nm) was used. The blue light was focused on to the back aperture of the microscope objective, producing a wide-field exposure around the recorded cell of 1 mW. The light power at the specimen was measured using an optical power meter PM100D (ThorLabs). The light output is controlled by a programmable pulse stimulator, Master-8 (AMPI Co. Israel) and the pClamp 10.2 software (AXON Instruments).

#### Activation of hM3Dq with CNO in AVP neurons

The modified human M3 muscarinic receptor hM3Dq (25) was expressed in AVP neurons by injecting a Cre-dependent virus containing hM3Dq (AAV-DIO-hM3Dq-mCherry) into the SON of mice bearing *Avp* promoter-driven Cre recombinase (*Avp^ires-^*^Cre+^ mice; see **Figure 1a**). The intracellular solution for current clamp recordings contained the following (in mM): 128 K gluconate, 10 KCl, 10 HEPES, 1 EGTA, 1 MgCl2, 0.3 CaCl2, 5 Na2ATP, 0.3 NaGTP, adjusted to pH 7.3 with KOH.

### Brain immunohistochemistry

Mice were terminally anesthetized with chloral hydrate (Sigma-Aldrich) and trans-cardially perfused with phosphate-buffered saline (PBS) followed by 10% neutral buffered formalin (Fisher Scientific). Brains were extracted, cryoprotected in 20% sucrose, sectioned coronally on a freezing sliding microtome (Leica Biosystems) at 40 μm thickness, and collected in two equal series. Brain sections were washed in 0.1 M PBS with Tween-20, pH 7.4 (PBST), blocked in 3% normal donkey serum/0.25% Triton X-100 in PBS for 1 h at room temperature and then incubated overnight at room temperature in blocking solution containing primary antiserum (rat anti-mCherry, Invitrogen M11217, 1:1,000; chicken anti-GFP, Life Technologies A10262, 1:1,000; rabbit anti-vasopressin, Sigma-Aldrich AB1565, 1:1,000; rabbit anti-TH, Millipore AB152, 1:1,000). The next morning, sections were extensively washed in PBS and then incubated in Alexa-fluor secondary antibody (1:1000) for 2 h at room temperature. After several washes in PBS, sections were incubated in DAPI solution (1 µg/ml in PBS) for 30 min. Then, sections were mounted on gelatin-coated slides and fluorescence images were captured using an Olympus VS120 slide scanner.

### Reagents

AVP, hydrocortisone and adrenaline were all from Sigma (Sigma-Aldrich, UK). The G_q_ inhibitor YM-254890 (39) was from Wako-Chem (Wake Pure Chemical Corp). The selective V1bR antagonist SSR149415 (nelivaptan; Serradeil-Le Gal, et al. (31)) was from Tocris (Bio-Techne Ltd, UK). The glucagon receptor antagonist LY2409021 (adomeglivant; Kazda, et al. (30)) was from MedKoo Biosciences (USA).

### Clamping studies in human participants

Clamping studies were conducted at Gentofte Hospital, University of Copenhagen. The studies were approved by the Scientific-Ethical Committee of the Capital Region of Denmark (registration no. H-D-2009-0078) and was conducted according to the principles of the Declaration of Helsinki (fifth revision, Edinburgh, 2000).

#### Comparison of copeptin in healthy subjects undergoing a hypoglycemic and euglycemic clamp

Samples from the “saline arm” from 10 male subjects enrolled in an ongoing, unpublished clinical trial (https://clinicaltrials.gov/ct2/show/NCT03954873) were used to compare copeptin secretion during euglycemia and hypoglycemia.

For the study, two cannulae were inserted bilaterally into the cubital veins for infusions and blood sampling, respectively. For the euglycemic study, participants were monitored during fasting glucose levels. For the hypoglycemic clamp, an intravenous insulin (Actrapid; Novo Nordisk, Bagsværd, Denmark) infusion was initiated at time 0 min to lower plasma glucose. Plasma glucose was measured bedside every 5 min and kept >2.2 mM. Arterialised venous blood was drawn at regular time intervals prior to and during insulin infusion.

#### Comparison of copeptin in subjects with T1DM and healthy controls

Samples from 20 male (n=10 control and n=10 T1DM patients) from the “saline arms” of two previously published studies (80, 81) were used to compare copeptin secretion during a hypoglycemic clamp between T1DM and control subjects. The samples from healthy individuals (Controls) were from Christensen, et al. (80). These 10 healthy male subjects were of age 23 ± 1 years, BMI 23 ± 0.5 kg/m^2^ and HbA_1c_ 5.5 ± 0.1%. The T1DM patient samples were from (81). These patients were; C-peptide negative, age 26 ± 1 years, BMI 24 ± 0.5 kg/m^2^, HbA_1c_ 7.3 ± 0.2%, positive islet cell and/or GAD-65 antibodies, treated with multiple doses of insulin (*N* = 9) or insulin pump (*N* = 1), without late diabetes complications, without hypoglycemia unawareness, and without residual β-cell function (i.e., C-peptide negative after a 5-g arginine stimulation test). For the study, a hypoglycemic clamp was conducted as outlined above.

#### Measurement of copeptin, glucagon and AVP in human plasma

Copeptin in human plasma was analysed using the KRYPTOR compact PLUS (Brahms Instruments, Thermo Fisher, DE). Glucagon was measured using human glucagon ELISA from Mercodia. Plasma AVP was measured using a human AVP ELISA kit (CSB-E09080h; Cusabio, China).

### Statistical tests of data

All data are reported as mean ± SEM. Unless otherwise stated, N refers to the number of mice. Statistical significance was defined as p < 0.05. All statistical tests were conducted in Prism8 (GraphPad Software, San Diego, CA, USA). For two groupings, a t test was conducted with the appropriate post hoc test. For more than two groupings, a one-way ANOVA was conducted (repeated measures, if appropriate). If data were separated by two treatments/factors, then a two-way ANOVA was conducted. A repeated measures (RM) two-way ANOVA was used (if appropriate), and a mixed-models ANOVA was used in the event of a repeated measures with missing data.

